# Activation of silent secondary metabolite gene clusters by nucleosome map-guided positioning of the synthetic transcription factor VPR-dCas9

**DOI:** 10.1101/2020.04.02.022053

**Authors:** Andreas Schüller, Lisa Wolansky, Harald Berger, Lena Studt, Agnieszka Gacek-Matthews, Michael Sulyok, Joseph Strauss

## Abstract

Current methods for forced expression of selected target genes are based on promoter exchange or on overexpressing native or hybrid transcriptional activators in which gene-specific DNA binding domains are coupled to strong activation domains. While these approaches are very useful for promoters with known or synthetically introduced transcription factor binding sites, they are not suitable to turn on genes in biosynthetic gene clusters which often lack pathway-specific activators. To expand the discovery toolbox, we designed a Cas9-based RNA guided synthetic transcription activation system for *Aspergillus nidulans* based on enzymatically disabled dCas9 fused to three consecutive activation domains (VPR-dCas9). Targeting two biosynthetic gene clusters involved in the production of secondary metabolites, we demonstrate the utility of the system. Especially in silent regions facultative heterochromatin and strictly positioned nucleosomes can constitute a relevant obstacle to the transcriptional machinery. To avoid this negative impact and to facilitate optimal positioning of RNA-guided VPR-dCas9 to our targeted promoters we have created a genome-wide nucleosome map to identify the cognate nucleosome-free-regions (NFRs). Based on these maps, different single-guide RNAs (sgRNA) were designed and tested for their targeting and activation potential. Our results demonstrate that the system can be used to activate silent BGCs in *A. nidulans*, partially to very high expression levels and also open the opportunity to stepwise turn on individual genes within a BGC that allows to decipher the correlated biosynthetic pathway.

## 1. Introduction

Filamentous fungi produce a plethora of metabolites and enzymes which are essential components of their response to environmental, nutritional or developmental signals. These metabolites can have beneficial but also detrimental effects on plant and animal health. In the agricultural sciences, fungal research is focused on understanding the biology and genetics of phytopathogens to combat diseases and to reduce economic losses caused by product deterioration and the accumulation of toxic secondary metabolites (SMs) (Brefort et al., 2009; Hollingsworth et al., 2008). Usually, multi-gene biosynthetic gene clusters (BGCs) are responsible for the production of these SMs. These clusters show similar structures at the genomic level always containing a core gene that encodes a backbone-generating enzyme that defines the substance class of the cluster product. These backbone or signature genes often code for polyketide synthases (PKSs), non-ribosomal peptide synthetases (NRPSs), prenyl-transferases or terpene cyclases. Apart from the core genes, several other protein classes can be involved that serve the purpose of applying modifications, regulating gene transcription or transport of the metabolite (Brakhage, 2013). Usually, the individual genes within a given BGC are transcriptionally co-regulated, not expressed by default and activated only in response to a “proprietary” expression signal. For most of these clusters, standard laboratory conditions do not generate this critical signal and consequently they are silent. (Bachleitner et al., 2019; Chujo and Scott, 2014; Connolly et al., 2013; Gacek-Matthews et al., 2016; Gacek and Strauss, 2012; Reyes-Dominguez et al., 2010; Studt et al., 2016). In about 50% of the cases the SM BGCs contain a gene that encodes for a transcriptional activator (Keller, 2019). In many cases the presence of these regulators is sufficient to orchestrate the production of the BGC product. However, the other half of the BGCs do not contain a known transcriptional activator or are regulated by regulators that are not located within the cluster boundaries and are involved in the regulation of primary as well as secondary metabolism. This complicates the search for conditions and activators of a specific cluster (Brakhage, 2013; Macheleidt et al., 2016; Then Bergh and Brakhage, 1998; Tilburn et al., 1995). The number of genes that are involved in the production of SMs can vary greatly. Some SMs need only a single gene like the *PKS8* in *Fusarium graminearum* (Westphal et al., 2018). Others need the coordinated involvement of few to many genes for the assembly of the BGC product like the asperthecin cluster of *Aspergillus nidulans* which comprises 3 genes (Szewczyk et al., 2008) or the sterigmatocystin cluster of *A. nidulans* which consists of 25 genes; (Brown et al., 1996). Genomes of filamentous fungi contain a large number and diversity of BGCs. It is well established that the repression of SM BGCs can comprise a multitude of global but also specific regulators that act on the maintenance of a repressive chromatin structure, influence signal transduction pathways or impact gene expression by post translational modifications/ processing of regulators. Therefore, the products of many silent SM BGCs remain elusive. For example, *A. nidulans* harbours 68 predicted BGCs, but only 20 metabolites are known and assigned to a specific BGC in this model fungus (Chiang et al., 2016; Macheleidt et al., 2016). The same is true for other fungi such as *Fusarium* spp. (Hansen et al., 2015; Niehaus et al., 2016a), *Penicillium* sp. (Nielsen et al., 2017) and *Sclerotinia sclerotiorum* (Graham-Taylor et al., 2020). Due to the recent revolution in sequencing techniques the complete genome sequence of over 700 different fungal species has been determined so far (https://mycocosm.jgi.doe.gov) and this theoretically represents an enormously rich resource for novel bioactive compounds (Nordberg et al., 2014). To lift this treasure, techniques to activate the transcription of these predominantly silent BGCs have been developed. They include promoter replacements *in cis*, overexpression of cluster-specific or global activators *in trans*, deletion of global regulators, interference with chromatin-based silencing or trans-expression of whole BGCs in a heterologous host and approaches based on the variation of cultivation conditions (i.e. “OSMAC”) (Bode et al., 2002; Brakhage and Schroeckh, 2011; Chiang et al., 2010; Clevenger et al., 2017; Connolly et al., 2013; Gacek-Matthews et al., 2016; Gerke et al., 2012; Gerke and Braus, 2014; Hansen et al., 2015; Keller, 2019; Lyu et al., 2020; Niehaus et al., 2016b; Strauss and Reyes-Dominguez, 2011; Studt et al., 2016; Westphal et al., 2019; Wiemann et al., 2013; Wiemann et al., 2018). Promoter exchanges seem only feasible with small predicted clusters or clusters that harbour a clear candidate of BGC-specific transcription factor (TF) that can be overexpressed and converted to an active state. For BGCs that do not contain a TF or contain a TF that requires specific post-translational modifications for activity, this approach will not be successful. Large promoter exchange approaches that aim for the overexpression of all genes of a cluster are also only applicable for a small number of target genes and require tedious work and selection marker recycling strategies which always change the nucleotide sequence at the locus of interest (e.g. Cre/loxP) (Aguiar et al., 2014). Another problem that emerges during promoter exchange studies is the integrity of the transformation locus. Fungi like *Aspergillus* spp. and *Fusarium* spp. have a high gene density (∼ 1 gene/ 3,000 bps (Bashyal et al., 2017; Cuomo et al., 2007; Galagan et al., 2005)) when compared to other eukaryotes like human (∼ 1 gene/ 150,000 bps (Ezkurdia et al., 2014; Piovesan et al., 2019)), *Mus musculus* (∼ 1 gene / 83,000 bps (Weitzman, 2002)), *Caenorhabditis elegans* (∼ 1 gene/ 5,000 bps (Consortium, 1998)), *Drosophila melanogaster* (∼ 1 gene/ 9,000 bps (Adams et al., 2000)), *Arabidopsis thaliana* (∼ 1 gene/ 5,000 bps (Arabidopsis Genome, 2000)) or *Oryzae sativa* (∼ 1 gene / 9,500 bps (Kawahara et al., 2013)). Thus, many control elements (promoters, terminators, binding motifs, etc.) in filamentous fungi like *A. nidulans* are in near vicinity to each other. Interfering with the integrity of the genomic locus could lead to unforeseen consequences on neighbouring genes, *cis*- or *trans-*regulatory elements and the chromatin structure at this locus. Apart from promoter replacement approaches, the construction of hybrid transactivators was tested for activation. By fusing the DNA-binding domain of the cluster-specific TF residing inside the (+)-Asperlin BGC with an activation domain of another transcriptional activator (i.e. AfoA), the cluster could be activated and the cognate product was identified (Grau et al., 2018). Again, this approach can only be successful if the BGC in question contains also a specific TF. Because of these complications and limitations of established techniques there are still hundreds of predicted fungal BGCs awaiting the characterisation of their cognate metabolite (Lyu et al., 2020). Other approaches that interfere with chromatin-based regulators, such as histone deacetylases (Shwab et al., 2007; Studt et al., 2013), histone methyl transferases (Chujo and Scott, 2014; Connolly et al., 2013; Studt et al., 2016),histone demethylases (Bachleitner et al., 2019; Gacek-Matthews et al., 2016), etc. may work for some clusters, but not for others. Furthermore, this approach cannot be used for targeting only a specific cluster but may have effects on many BGCs simultaneously and also can impact primary metabolism and developmental processes as well.

Here, we present a new method to activate silent BGCs in fungi which is based on the expression and precise targeting of a synthetic activator composed of an enzymatically disabled “dead” Cas9 (dCas9) fused to three consecutively arranged strong activation domains termed V-P-R. The whole dCas9-VPR system was already described to function in human cell lines by Chavez et al. (2015). In the tripartite activation domain “V” represents 4 copies of the VP16 domain of the Herpes Simplex Virus (i.e. VP64), “P” stands for a part of the p65 domain of the human transcription factor nf-κB and “R” represents a segment of the Rta transactivator of the Epstein-Barr virus. Each of these activator domains have been extensively studied and shown to individually enhance or facilitate gene activation by engaging in different protein interactions within the transcriptional machinery (Chang et al., 2005; Hall and Struhl, 2002; Hardwick et al., 1992; Lecoq et al., 2017; Lee and Hahn, 1995; Schmitz and Baeuerle, 1991). We centred our approach on a dCas9-based gene activation system because with this, the activator can be targeted at one or more genomic locations easily and simultaneously simply by expressing the right combination of guide RNAs. Moreover, the functionality of the conventional CRISPR/Cas9 system for genetic manipulation of a single site or multiple loci (multiplex sgRNAs) is firmly established in *A. nidulans* and other filamentous fungi as reviewed by Song et al. (Nodvig et al., 2018; Song et al., 2019). We reasoned that VPR-dCas9 can be expressed in the host and guided by co-expressed sgRNAs to any accessible locus in the genome. By expression of one or more sgRNAs in the *VPR-dCas9*-containing strain one or more individual genes in different combinations can be targeted and activated in a predicted BGC. This provides the unique opportunity to stepwise activate different genes within a target BGC eventually leading to the discovery of a novel metabolite and its intermediates. Here, we detail the methodology and provide proof-of-concept for functionality of the system by targeting the known – but silent – monodictyphenone (*mdp*) cluster as well as two genes in a predicted cluster for which the cognate product is not known.

## 2. Materials and Methods

### 2.1. Plasmid generation

Plasmids were constructed by means of DNA restriction digests, polymerase chain reactions (PCRs) and subsequent ligation by yeast recombinational cloning (YRC) (Schumacher, 2012). All oligos were acquired from Sigma-Aldrich (“DNA Oligos in Tubes”). For YRC, overlap extension PCRs were used for creating fragments with homologous overhangs (≥20 nucleotides). The backbone plasmid was pRS426 (Christianson et al., 1992) and the yeast strain for transformation was the uracil-auxotrophic strain FGSC 9721 (FY 834) (Winston et al., 1995). 5 µL of each PCR fragment was used for YRC. After YRC, the purified plasmids (New England Biolabs: Monarch^®^ Plasmid Miniprep Kit; Art. No.: T1010S) were verified by sequencing (“Ready2Run” by LGC Genomics GmbH). All plasmids that have been generated and used in that study are summarised Table 1. All fragments and the corresponding primers are listed in Table S 1.

**Table 1.** *Plasmids. Plasmid names, the most important elements and the backbone plasmid are shown. Consecutive numbers in the plasmid name (i.e.: m1-m4 and A1-A2) indicate different sgRNAs. Targeted genes are shown under elements in parenthesis. Elements comprise the human estrogen Receptor (*hER*), the activator* VPR-dCas9, the selection markers (i.e.: *argB* or *pyroA*) and the different sgRNA cassettes.

| Plasmid | Elements | Backbone |
| --- | --- | --- |
| pVPR4 | *hER*, *VPR-dCas9*, *argB* | pRS426 |
| psgRNA | sgRNA scaffold, *pyroA* | pRS426 |
| psgRNA-m1 | sgRNA-m1 (*mdpE*), *pyroA* | psgRNA |
| psgRNA-m2 | sgRNA-m2 (*mdpE*), *pyroA* | psgRNA |
| psgRNA-m3 | sgRNA-m3 (*mdpE*), *pyroA* | psgRNA |
| psgRNA-m4 | sgRNA-m4 (*mdpE*), *pyroA* | psgRNA |
| psgRNA-A1 | sgRNA-A1 (AN8506), *pyroA* | psgRNA |
| psgRNA-A2 | sgRNA-A2 (AN8507), *pyroA* | psgRNA |

<u>For generation of pVPR4</u> (for plasmid map see Figure 1A; The number indicates the laboratory internal plasmid version), the backbone vector pRS426 was digested with EcoRI and XhoI over night at 37°C in 2x Tango buffer (ThermoFisher Scientific: Art. No.: ER0271, ER0691) and directly used for YRC. Together with the digest, following DNA fragments were used for transformation of yeast strain FY834 (For Primer pairs see Supplementary Table S 1). The gene of the human estrogen receptor alpha (*hER*) was amplified by primer pair hER-F/R (template source: phERpyr4 (Pachlinger et al., 2005), which is under the control of the constitutive *coxA* promoter (Sarkari et al., 2017) which was amplified by primer pair P.coxA-1-F/R and P.coxA-2-F/R (template source: genomic DNA from *Aspergillus niger* wild-type isolate) and the terminator region of the *tef1* orthologue of *A. niger* (An18g04840) which was amplified by primer pair T.tef1-1-F/R (template source: genomic DNA from *A. niger* wildtype isolate). The *VPR-dCas9* fusion was amplified from two different sources. *VPR* was amplified by primer pair VPR-F/R (template source: pWalium20-10XUAS-3XFLAG-dCas9-VPR (Addgene No.: # 78897)(Lin et al., 2015)) while *dCas9* was amplified by primer pair dCas9-1-F/R and dCas9-2-F/R (template source: pCAG-BirA*-dCas9-eGFP which was kindly provided by the Leonardt laboratory (Schmidtmann et al., 2016). The estrogen response elements which represent the promoter of the *VPR-dCas9* fusion gene were amplified by primer pair P.ERE-F/R (template source: pJW52_[3xERE-RS-P(nirA)]-[hER-trpC]-[pyro3/4] (Bödi, 2008). The terminator of *VPR-dCas9* is from the *glucanase* gene of *Botrytis cinerea* and was amplified by primer pairs T.gluc-F/R (template source: pNAH-P*oliC*::*bcltf3*-*gfp* (Brandhoff et al., 2017)). For selection of transformants, the whole *argB* gene (including promoter and terminator region) from *A. nidulans* was used as auxotrophy marker which was amplified by primer pair argB-F/R (template source: gDNA of *A. nidulans* FGSC A4).

**Figure 1:**
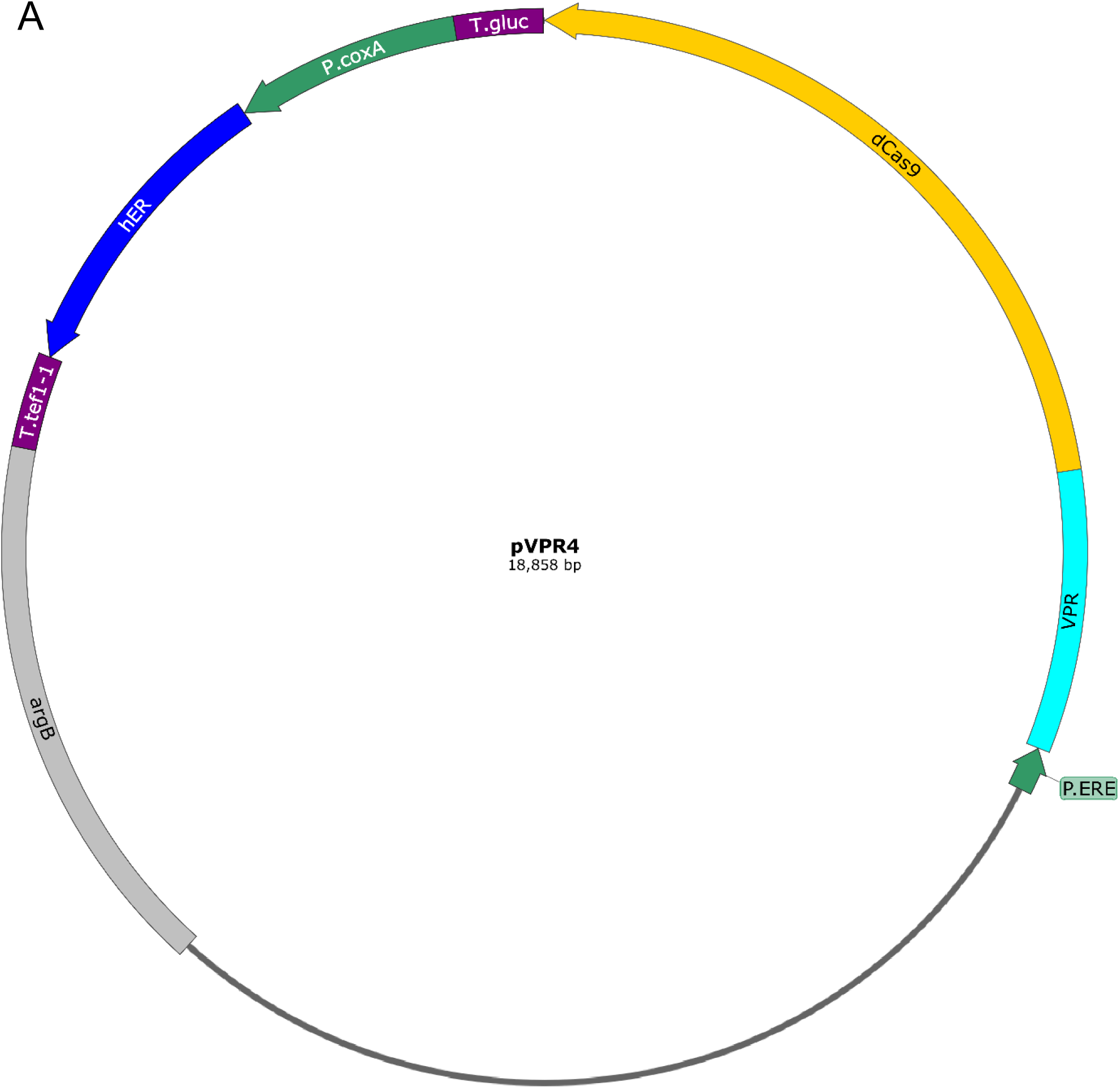
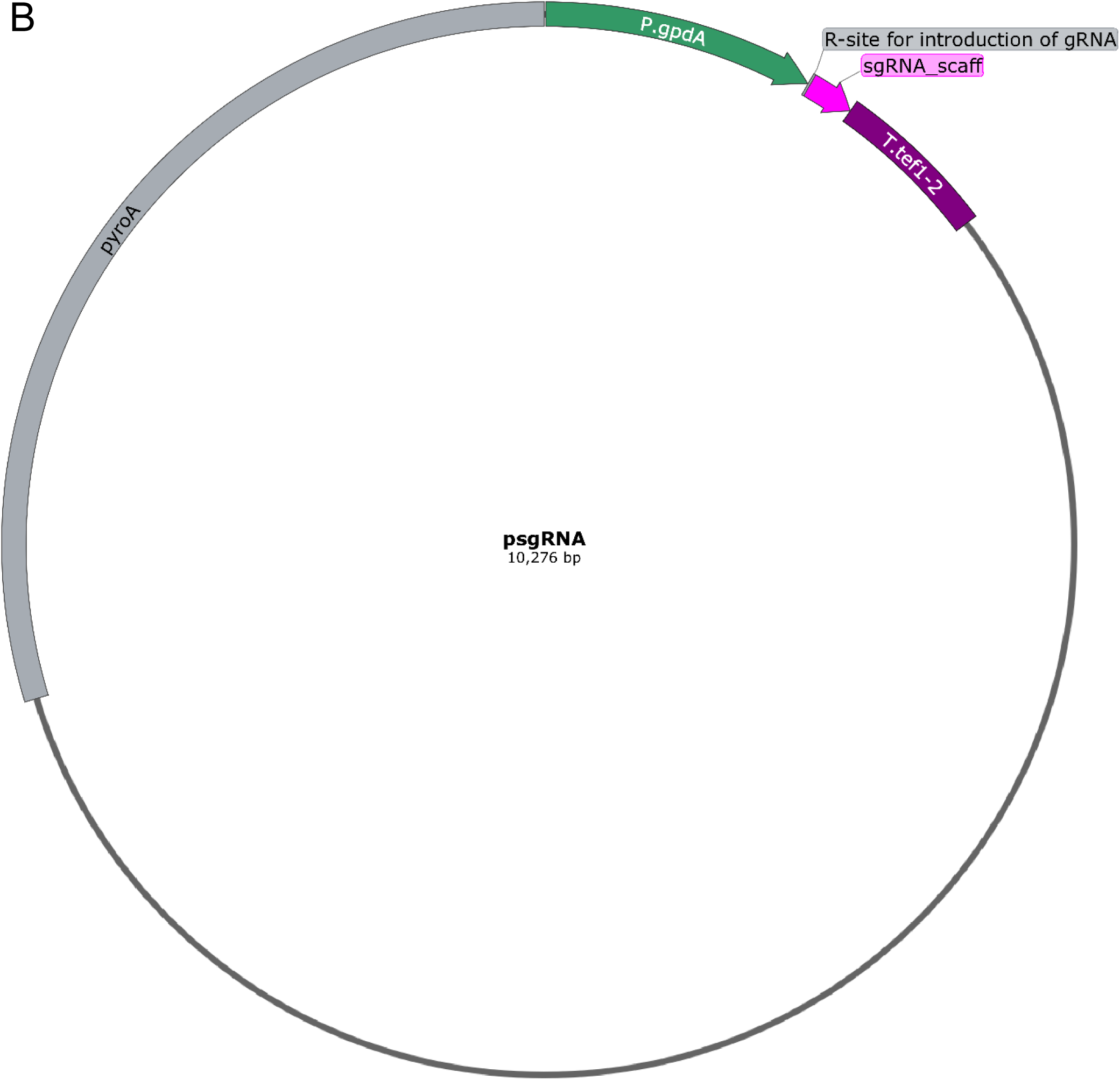
Schematic drawing of plasmids encoding the components of the VPR-dCas9 activation system. Selection markers for fungal transformation are depicted in grey, promoters in green and terminators in purple. **A:** Plasmid VPR4 carries the gene for the activator *VPR-dCas9 (cyan, yellow)* which has a synthetic promoter containing estrogen response elements (EREs) as binding motif for the human estrogen receptor (hER). The terminator of *VPR-dCas9* is derived from the *glucanase* gene of *Botrytis cinera*. The *hER* gene (blue) is transcribed constitutively by the *coxA* promoter from *Aspergillus niger* and termination is controlled by the terminator region of the *tef1* orthologue of *A. niger*. The produced hER protein remains transcriptionally inactive unless it is converted to an active form (i.e. homodimer formation) by the addition of estrogens (i.e. DES). The selection marker *argB* is originated from from *A. nidulans*. **B:** Plasmid sgRNA is the parent vector of all sgRNA carrying plasmids. It carriers the sgRNA scaffold (magenta) which consists of the tracrRNA and HDV ribozyme. Directly upstream of the tracrRNA sequence an Eco91I restriction site is introduced for insertion of the specific gRNA-sequences together with the hammerhead ribozyme. After insertion, both elements combined form the functional sgRNA cassette which releases the functional sgRNA upon transcription. The sgRNA cassette is constitutively expressed by the *gpdA* promoter of A. fumigatus and terminated by the terminator of the *tef1* gene of *A. fumigatus*. The selection marker pyroA originates from *A. fumigatus.* Detailed information about the plasmids and their construction can be retrieved from section 2.1.

For the generation of psgRNA (Figure 1B) and subsequent plasmids (i.e. psgRNA carrying a sgRNA), pRS426 was digested with NdeI (ThermoFisher Scientific: Art. No.: ER0581). Together with the digest, following DNA fragments were used for transformation of FY834 (see Supplementary Table S 1 for information about primers). The missing part of the *URA3* marker (disrupted by NdeI) was amplified by primer pair URA3-F/R as well as the *2µ-ori* which was amplified by primer pair 2µ-ori-F/R (template source: pRS426). The plasmid furthermore contains the sgRNA cassette which was amplified by primer pair sgRNAscaff-F/R (template source: pFC334 (Nodvig et al., 2015), which was kindly provided by the Mortensen laboratory). The promoter for the sgRNA-cassette is from the *gpdA* gene and was amplified by primer pair P.gpdA-F/R (template source: gDNA from *Aspergillus fumigatus* wild-type isolate). The terminator was from the *tef1* gene and was amplified by primer pair tef1-2-F/R (template source: gDNA of *A. fumigatus* wild-type isolate). The *pyroA* gene (including promoter and terminator) from *A. fumigatus* was used as auxotrophy marker for selection of transformants and was amplified by primer pair pyroA-F/R (template source: gDNA of *A. fumigatus* wild-type isolate)

For the insertion of sgRNAs into psgRNA (i.e. generation of plasmids psgRNA-m1 to m4 and psgRNA-A1 and A2), the plasmid was digested by Eco91I (ThermoFischer Scientific: Art. No.: ER0392). The digested plasmid was then used together with 2 µL of the respective annealed gRNA oligonucleotides for transformation of FY834. For annealing of the forward and reverse gRNA oligonucleotide (Supplementary Table S 2), both were dissolved in annealing buffer (final concentrations: 10 mM Tris, pH7.5, 50 mM NaCl, 1mM EDTA, 100 µM of each oligonucleotide). The DNA was subsequently denaturised by heating the mix up to 95°C for 5 minutes in a Thermocycler (Biometra, Thermocycler, Art. No.: T3000-48) followed by an annealing step by cooling the mix down to 25 °C at 0.03 °C/sec (Protocol by Sigma-Aldrich (Website: Sigma-Aldrich,, Access Date: 21.01.2020)).

### 2.2. Strain generation and molecular methods

#### 2.2.1. Strain generation

Strains used in this study are depicted in Table 2. All strains had the *nkuAΔ* background to reduce non-homologous end joining and facilitate homologous recombination events (Nayak et al., 2006). Strains were transformed by using protoplast transformation (Tilburn et al., 1983). All transformations were conducted with circular plasmids. For generation of strain VPR4, strain A1153 was transformed with pVPR4.The *in situ* integration (i.e. *argB*). After single spore isolation, the presence of all relevant features of pVPR4 was verified by PCR. Primer pairs that were used for the verification of strain VPR4 and the results can be obtained from supplementary Table S 1 and Figure S 1, respectively. A correct transformant was then transformed with plasmid psgRNA containing a sgRNA. In case of multiple sgRNA-carrying strains (i.e. VPR4-mAll, VPR4-m1&m2, VPR4-m2&m3) plasmids psgRNA-m1 to m4 were transformed simultaneously and the uptake of different combinations was verified by PCR. In case of strain VPR4-A1 and A2 either psgRNA-A1 or psgRNA-A2 was transformed into VPR4. For sgRNA-carrying strains, the presence of the specific sgRNA was verified by PCR as well. Information about primer pairs and results of the PCRs can be obtained from supplementary Table S 1 and Figure S 2, respectively. DNA was extracted according to (Cenis, 1992) with following deviation. Prior to extraction, the fungus was grown on solid Aspergillus minimal media (AMM; (Barratt et al., 1965)) and subsequently a thin layer of mycelia was transferred into a Lysing matrix A tube (MP Biomedicals; Art. No.: SKU 116910100). The material was then homogenised in a FastPrep-24™ (MP Biomedicals, Art. No.: SKU 116004500) at 6 m*s^-1^ for 20 seconds in Cenis lysis buffer. DNA was used for fragment amplification for YRC (Q5^®^ polymerase (New England Biolabs; Art. No.: M0491L) and for diagnostic PCR reactions (GoTaq^®^ Green Master Mix (Promega; Art. No.:M7845). PCR was performed according to the respective manufacturer’s instruction. Annealing temperature was set to 58 °C for Q5^®^ amplifications and to 60°C for GoTaq^®^ amplifications. For RNA isolation, freeze-dried mycelium was ground to a fine powder using mortar and pestle under liquid nitrogen and RNA was extracted with TRIzol™ Reagent (Invitrogen; Art. No.: 15596026) according to the manual. RNA content was measured using a NanoDrop™ spectrophotometer (Thermo Scientific™; Art. No.: ND-2000C). For cDNA synthesis, 2 µg total RNA was treated with DNaseI (Thermo Scientific™; Art. No.: EN0521). Complete removal of any residual DNA was then verified by PCR with the primer pair for the gene *actA* (AN6542), actA-control (Supplementary Table S 1). Next, 1 µg of DNA-free sample was taken for cDNA synthesis (Biorad; iScript™ cDNA Synthesis Kit, Art. No.: 1708891). Successful cDNA synthesis was again verified using the primer pair actA-control (Supplementary Table S 1).

**Table 2:**
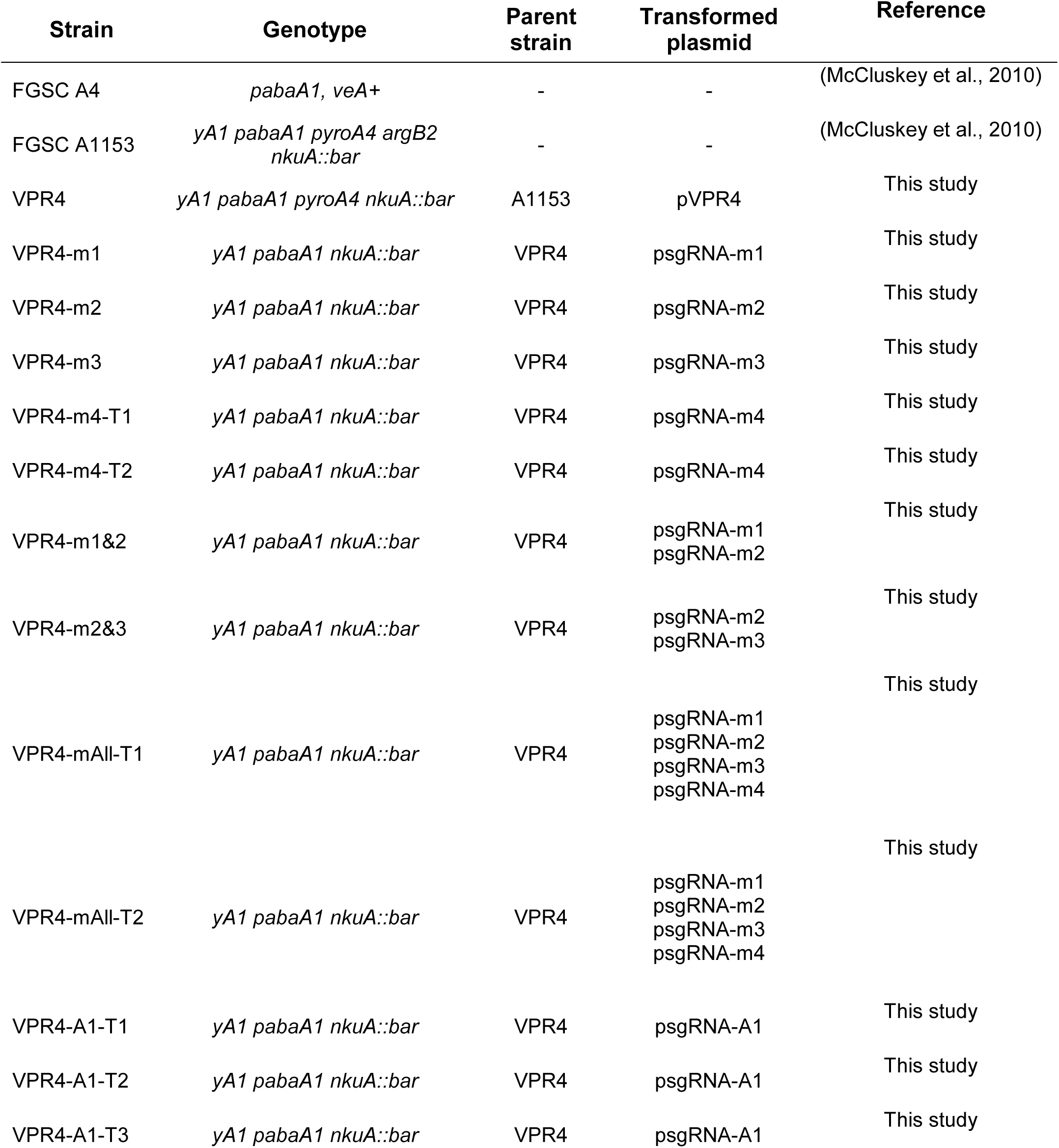
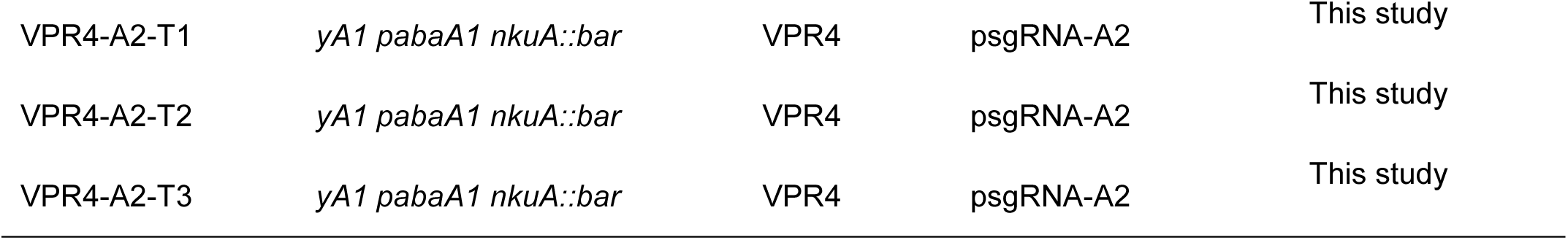
<u>*Aspergillus nidulans* strains</u> used during this study. All strains that are used for activation experiments are descendants of Strain FGSC A1153. The suffixes T1 / T2 / T3 in column “Strain” refer to the different independent transformants per construct.

| Strain | Genotype | Parent strain | Transformed plasmid | Reference |
| --- | --- | --- | --- | --- |
| FGSC A4 | *pabaA1*, *veA*<sup>+</sup> | - | - | (McCluskey et al., 2010) |
| FGSC A1153 | *yA1 pabaA1 pyroA4 argB2 nkuA::bar* | - | - | (McCluskey et al., 2010) |
| VPR4 | *yA1 pabaA1 pyroA4 nkuA::bar* | A1153 | pVPR4 | This study |
| VPR4-m1 | *yA1 pabaA1 nkuA::bar* | VPR4 | psgRNA-m1 | This study |
| VPR4-m2 | *yA1 pabaA1 nkuA::bar* | VPR4 | psgRNA-m2 | This study |
| VPR4-m3 | *yA1 pabaA1 nkuA::bar* | VPR4 | psgRNA-m3 | This study |
| VPR4-m4-T1 | *yA1 pabaA1 nkuA::bar* | VPR4 | psgRNA-m4 | This study |
| VPR4-m4-T2 | *yA1 pabaA1 nkuA::bar* | VPR4 | psgRNA-m4 | This study |
| VPR4-m1&2 | *yA1 pabaA1 nkuA::bar* | VPR4 | psgRNA-m1<br>psgRNA-m2 | This study |
| VPR4-m2&3 | *yA1 pabaA1 nkuA::bar* | VPR4 | psgRNA-m2<br>psgRNA-m3 | This study |
| VPR4-mAll-T1 | *yA1 pabaA1 nkuA::bar* | VPR4 | psgRNA-m1<br>psgRNA-m2<br>psgRNA-m3<br>psgRNA-m4 | This study |
| VPR4-mAll-T2 | *yA1 pabaA1 nkuA::bar* | VPR4 | psgRNA-m1<br>psgRNA-m2<br>psgRNA-m3<br>psgRNA-m4 | This study |
| VPR4-A1-T1 | *yA1 pabaA1 nkuA::bar* | VPR4 | psgRNA-A1 | This study |
| VPR4-A1-T2 | *yA1 pabaA1 nkuA::bar* | VPR4 | psgRNA-A1 | This study |
| VPR4-A1-T3 | *yA1 pabaA1 nkuA::bar* | VPR4 | psgRNA-A1 | This study |
| VPR4-A2-T1 | *yA1 pabaA1 nkuA::bar* | VPR4 | psgRNA-A2 | This study |
| VPR4-A2-T2 | *yA1 pabaA1 nkuA::bar* | VPR4 | psgRNA-A2 | This study |
| VPR4-A2-T3 | *yA1 pabaA1 nkuA::bar* | VPR4 | psgRNA-A2 | This study |

#### 2.2.2. Reverse Transcriptase quantitative polymerase chain reaction (RT-qPCR) analysis

The cDNA was diluted 1/5 with dH_2_O and 2.5 µL were used for qPCR analysis (10 µL total volume; BioRad, iTaq Universal SYBR Green Supermix; Art. No.: 172-5124) in a Biorad CFX384 C1000 Touch Thermal cycler. The qPCRs were done in technical duplicates. Only primer pairs with an efficiency between 90 and 110% were considered for qPCR analysis. The genes of interest were compared to the housekeeping genes *actA* (AN6542) and *benA* (AN1182) and the relative fold change in expression was calculated according to the 2^-ΔΔCt^ method (Livak and Schmittgen, 2001). The qPCR program was run as following: 95°C (3 min) – 40 repetitions of [95°C (10 s) – 60°C or 64°C (10 s) – 72°C (30 s)]-95°C (10 s) – Melting curve [65°C -95°C at 0.5°C intervals and 5 s hold at each interval]. All qPCR-primer pairs were run at an annealing temperature of 60 °C apart from primers for *mdpG* which were run at 64 °C. A complete list of qPCR Primers can be found in supplementary Table S 1.

### 2.3. Nucleosome mapping

*A. nidulans* (Strain FGSC A4) was grown in shaking flasks for 15 or 48 hours in AMM supplemented with 1% glucose, 100 mM sodium nitrate and para-Aminobenzoic acid (paba). Before harvest, mycelia were fixed with 1% formaldehyde (Roth; Art. No.: 4979) for 15 minutes and fixation was stopped with 1M glycine (Roth; Art. No.: 3790). Mycelia were filtered and shock frozen in liquid nitrogen prior to grinding using a mortar and pestle.

To fragment the DNA, mycelia were suspended in 2 mL MNase digestion buffer (50 mM Hepes pH 7.5 (Sigma; Art. No.: 3375), 50 mM NaCl (CarlRoth; Art. No.: 3957), 5 mM MgCl_2_ (CarlRoth; Art. No.: KK36.1), 1 mM CaCl_2_ (CarlRoth; Art. No.: A119.1), 1× Proteinase inhibitor mix (Sigma; Art. No.: P8215) and 1 mM phenylmethylsulfonyl fluoride (PMSF, Sigma; Art. No.: 6367) and 300 µL aliquots of suspension were treated with 0.4 U micrococcal nuclease (MNase, Sigma ; Art. No.: N3755) at 37 °C for 6 min. The MNase reaction was stopped by adding 300 µl stop buffer 2 (50 mM Hepes pH 7.5, 255 mM NaCl, 40 mM EDTA (CarlRoth; Art. No.: 8040), 2% Triton X-100 (Sigma; Art. No.: 8787), 0.2% sodium deoxycholate (Sigma; Art. No.: D6750). Fixation was reversed by incubation for 15 min at 65°C, the supernatant was purified using a PCR purification kit (Qiagen; Art. No.: 28181) and samples were electrophoresed on a 1.5 per cent agarose gel to check the MNase digestion. Sequencing library preparation and sequencing was performed at Vienna Biocenter Core Facilities. Illumina libraries were prepared using Nextera XT DNA Library Preparation Kit and paired end 50 bp sequencing was done on an Illumina Hi-Seq 2000 system. Obtained fastq files were mapped on *A. nidulans* FGSC A4 genome assembly using BWA (version 0.7.17) and coverage per base pair was calculated using samtools (version 1.9) and bedtools (version 2.28.0) to receive a representation of the genome wide nucleosome occupancy.

### 2.4. Guide RNA design

The region of interest was analysed for the occurrence of the protospacer adjacent motifs (PAMs; i.e. NGG) and possible protospacer sequences (i.e. target specific sequence of sgRNA) were subjected to a full genome blast against the genome of *A. nidulans* to select the protospacers that are closest to the anticipated locus with least possible off-targets. The borders of nucleosome free regions were designated as the region between a 75 base pair offset from the adjacent dyad axis (i.e. peak maximum) (see Figure 4 and Figure 6). In case of the activation of *mdpE*, sgRNA sequences upstream as well as downstream of the nucleosome free region were chosen additionally. Two sgRNAs were positioned close to the predicted transcription start point (TSP) of *mdpE*. Sequences of sgRNAs can be retrieved from supplementary Table S 2. The positioning of the sgRNAs within the genome is shown in Figure 4 and Figure 6.

**Figure 2:**
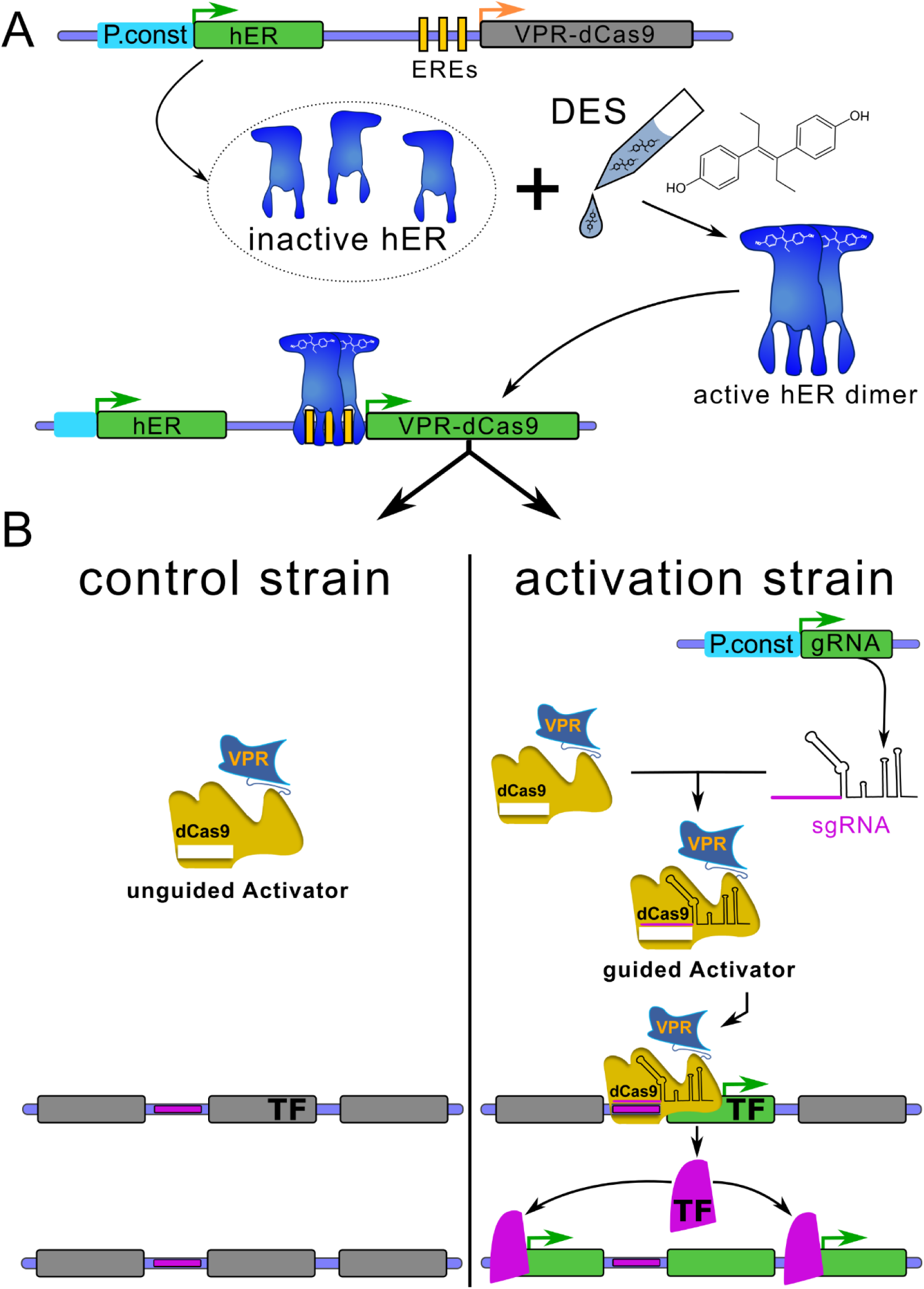
VPR-dCas9 activation and experimental design. Green or grey rectangulars represent expressed or not expressed genes respectively. Green arrows indicate active promoters while orange arrows represent promoters poised for activation. **A:** All Strains carry plasmid VPR4 (Control as well as activation strains). pVPR4 contains the constitutively expressed (P.const; cyan) *human estrogen receptor (hER)* which is inactive in absence of the inducer. Upon induction with DES, hER is activated and able to bind to estrogen response elements (EREs; vertical yellow bars) which are present in the engineered *VPR-dCas9* gene promoter. Upon binding, hER facilitates transcription of the gene that codes for VPR-dCas9 **B:** In the control strain (only pVPR4) sgRNAs are not expressed and thus VPR-dCas9 is present in the cell but not targeted to the designated region of interest (horizontal violet bars). The activation strains (pVPR4 and one or more psgRNA with functional sgRNA cassette), constitutively express sgRNA(s) (target sequence in violet) which guide VPR-dCas9 to the region of interest (horizontal violet bars) and facilitate transcription of the targeted gene (i.e. TF).Hence, the activation strain should experience an elevated expression of the targeted gene compared to the control strain, which can only be attributed to the presence of the sgRNA. In one application, the targeted gene could be a transcription factor gene (TF) of a silent BGC and forced expression of this TF gene could subsequently lead to the upregulation of the whole cluster as shown in the figure. In another scenario, several sgRNAs could be targeted to many genes within a predicted cluster and the cognate metabolite(s) could be identified subsequently.

**Figure 3:**
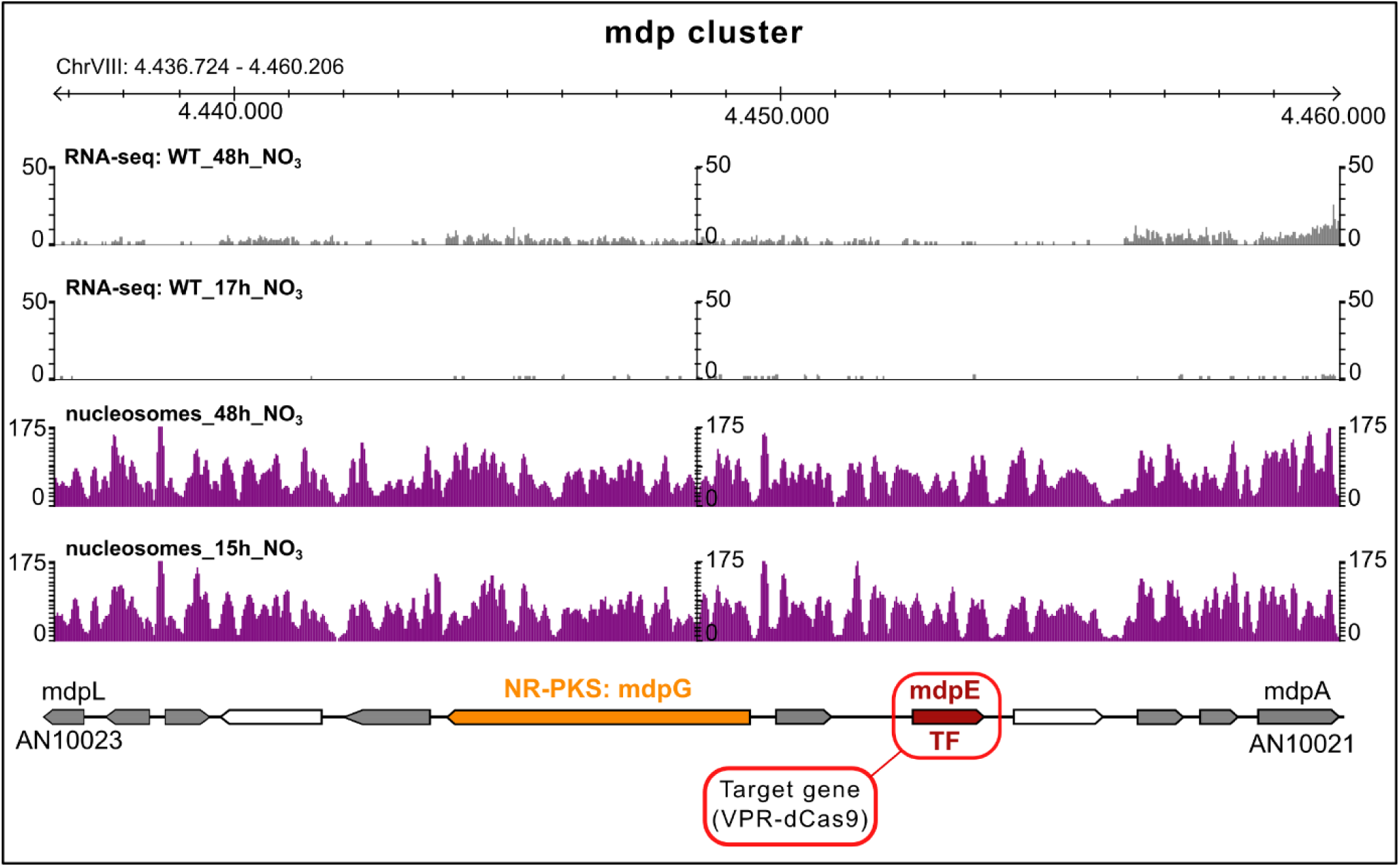
Overview of the monodictyphenone cluster of *A. nidulans*. The section spans the whole cluster. Apart from gene annotations (arrows), nucleosome positioning as well as the mRNA profile at nutrient rich (15/17h) and nutrient depleted (48h) conditions are depicted in purple and grey histograms respectively. White arrows are genes that are not necessary for the synthesis of the BGC products. Grey, orange and ruby arrows are genes involved in the monodictyphenone biosynthesis. The orange arrow is the backbone gene of the cluster which encodes for the nonreduced polyketide synthase MdpG. The ruby arrow is the gene that encodes for the pathway specific transcription factor MdpE which is the target for the activation by VPR-dCas9.

**Figure 4:**
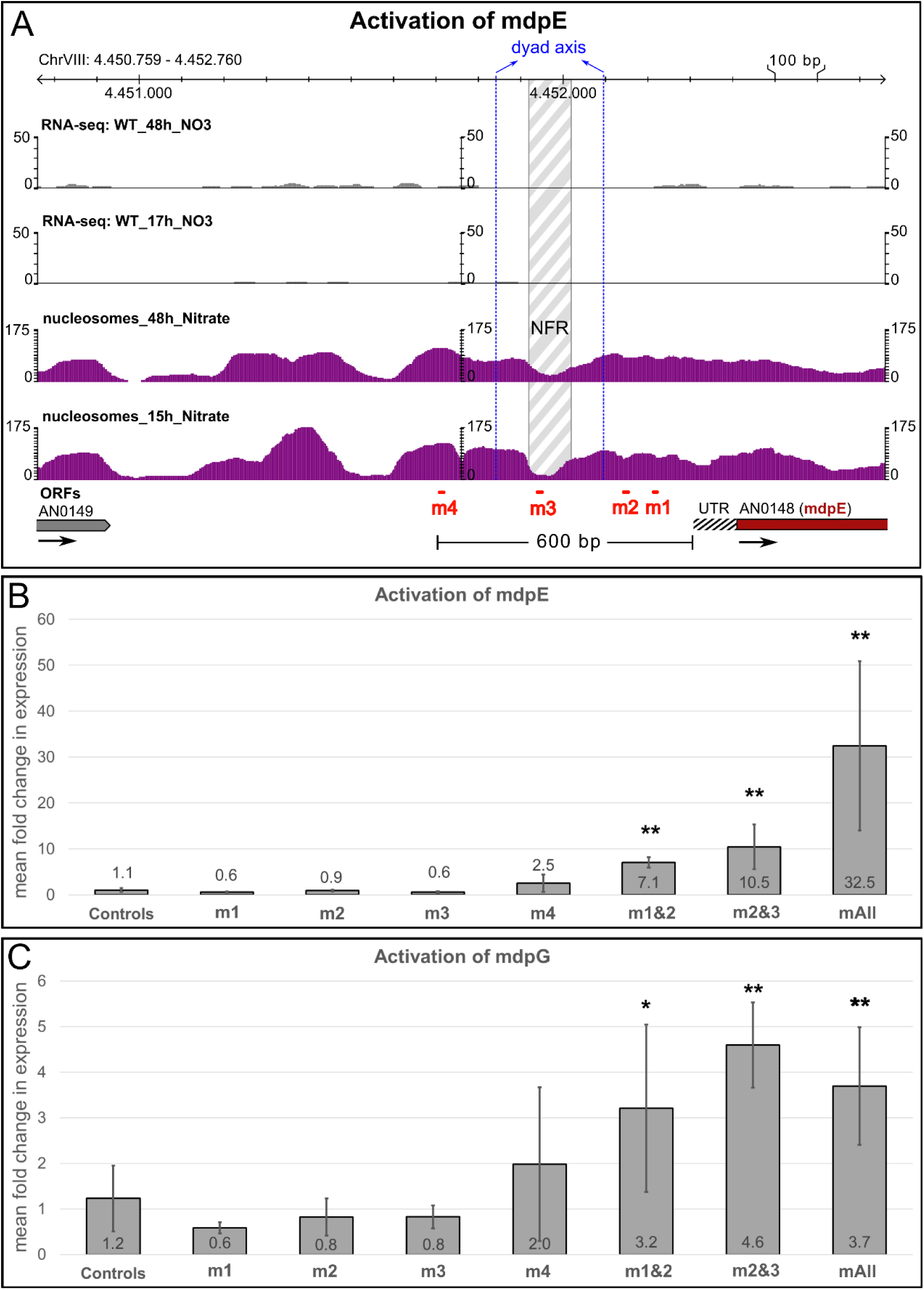
Targeted activation of the transcription factor gene *mdpE* via VPR-dCas9 and MdpE mediated activation of the backbone gene *mdpG*. **A:** Promoter region of the target gene *mdpE* together with RNA-sequencing data (grey histograms) and nucleosome positioning maps (purple histograms) under nutrient rich (15/17h) and nutrient depleted (48h) conditions. The nucleosome free region (NFR) is shown in grey white pattern which is delimited by the nucleosomal borders (set to an offset of 75 base pairs proximal to the dyad axis of the neighboring nucleosomes). The dyad axis are shown in vertical blue dashed lines and were set at the peak of the nucleosome histogram. Positions of sgRNAs are depicted in red (m1-m4). **B:** Expression of the gene *mdpE* in the controls (only pVPR4) and strains that additionally carry one or more sgRNAs (m1 | m2 | m3 | m4 | m1&2 | m2&3 | mAll) that target the promoter region of the gene *mdpE*. **C:** Expression of the backbone gene *mdpG* in the same samples as shown in Figure 4B which is caused by the activation of mdpE. Cultures were done in triplicates, qPCR was done in technical duplicates. A student’s t test was used to verify the significance of the activation over the controls (i.e. VPR4) (*p<0.05; **p < 0.01).

**Figure 5:**
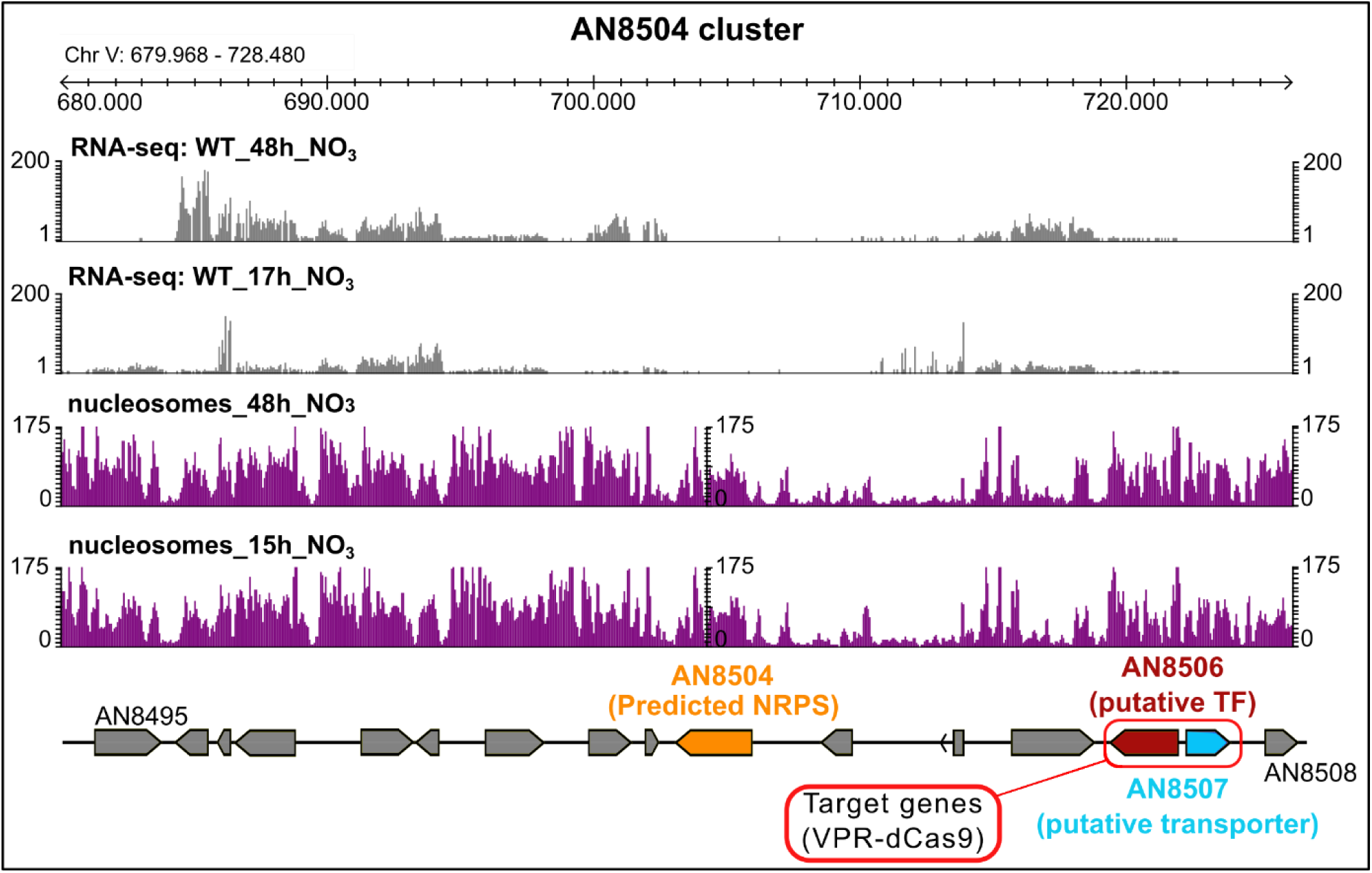
Overview of the predicted cluster AN8504 of *A. nidulans*. The cluster (as predicted by antiSMASH) begins at gene AN8495 and ends at AN8508. Nucleosome positioning as well as the mRNA profile at nutrient rich (15/17h) and nutrient depleted (48h) conditions is depicted in purple and grey histograms respectively. The backbone gene (AN8504; orange arrow) shows no transcription at either condition. Two sgRNAs were designed that target VPR-dCas9 to the bidirectional promoter that regulates the genes AN8506 and AN8507 (red frame). AN8506 (ruby arrow) is the putative transcription factor and AN8507 (cyan arrow) is the putative membrane protein. AN8506 shows a low basal transcription level (comparable to 0.32 times of the expression level of the housekeeping gene *benA*) while AN8507 shows no reads at either condition.

**Figure 6:**
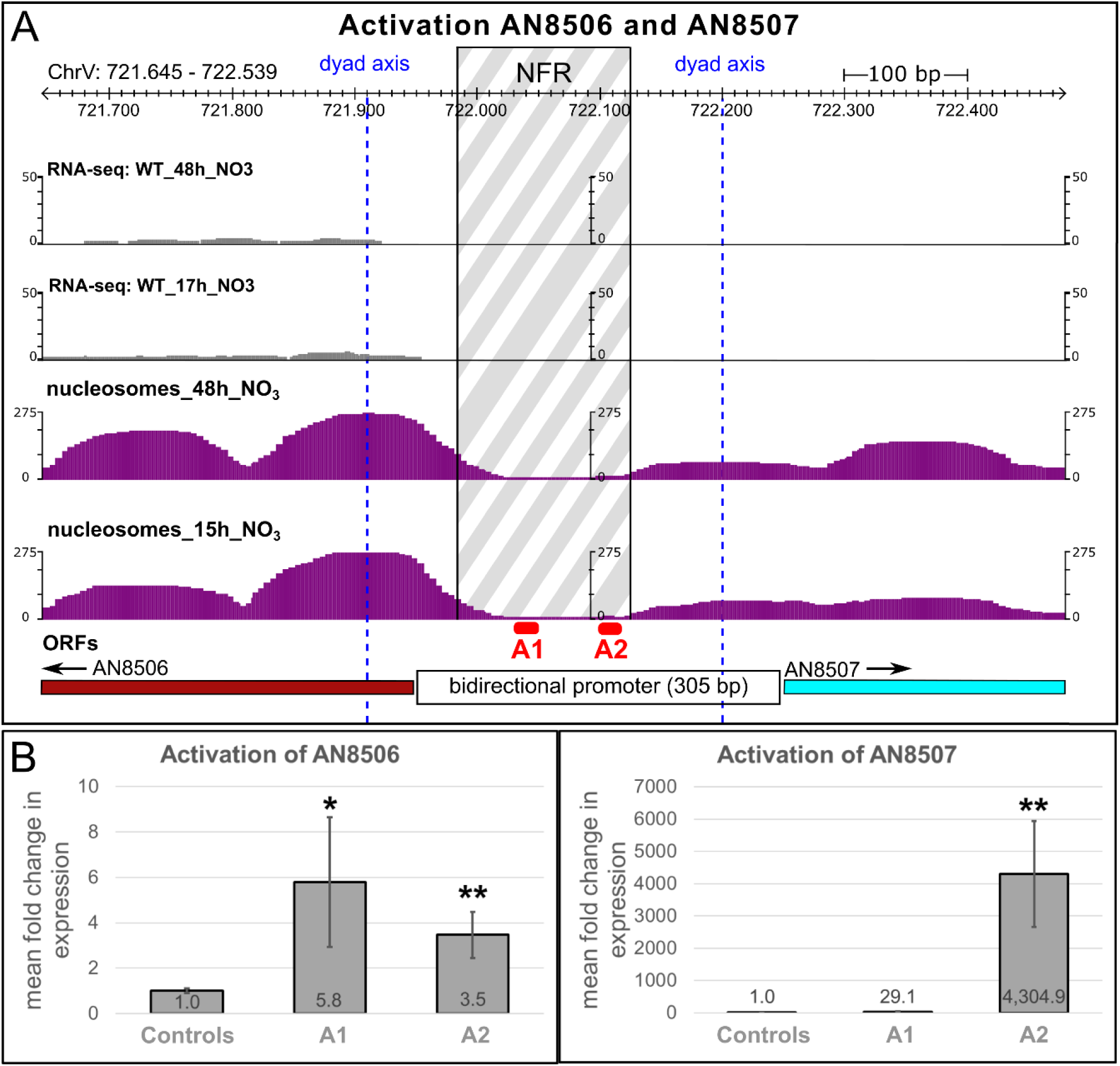
Targeting of VPR-dCas9 to a bidirectional promoter and activation of its underlying genes AN8506 and AN8507. **A**: The section shown comprises the begin of both genes AN8506 and AN8507 which share a bidirectional promoter of around 305 base pairs. The histograms show the mRNA profile (grey) and the nucleosome positioning (purple) under nutrient rich (15/17h) and nutrient depleted (48h) conditions. The nucleosome free region (NFR) is shown in grey white pattern which is delimited by the nucleosomal borders (set to an offset of 75 base pairs proximal to the dyad axis of the neighboring nucleosomes). The dyad axis are shown in vertical blue dashed lines and were set at the peak of the nucleosome histogram. Positions of sgRNAs are depicted in red (A1 and A2). **B:** Expression data of the activation experiment: Controls: VPR4 (no sgRNA), A1: VPR4-A1-T1 to T3 | A2: VPR4-A2-T1 to T3. Three independent transformants per sgRNA (T1, T2, T3). Cultures were executed in triplicates. qPCR was performed in technical duplicates. Reproducibility was verified by two independent conductions of the experiment. The significance of the activation effect was verified by a student’s t test (* p<0.05; **p < 0.01)

### 2.5. Media and growth conditions for activation experiments

During activation experiments, controls as well as activation strains were all grown and treated under identical conditions. 4*10^6^ conidia per mL AMM media were inoculated in 60 mL AMM liquid media and 10 mM nitrate as nitrogen source. The cells were grown for 40 hours at 37°C and 180 rpm on orbital shaker platforms before adding the inducer to all flasks (i.e.: samples as well as controls (=VPR4)). The Inducer mix contained 75 nM diethylstilbestrol (DES) in ethanol absolute (glass flask: Sigma; EMPARTA^®^ ACS: Art. No.: 1070172511) (final inducer concentration: 33 pM DES). After induction, cultures were incubated for further 8 hours. Subsequently 1 mL of supernatant was taken for SM analysis as described in section 2.7. Mycelia was harvested by filtering through miracloth fabric (Millipore; Art. No.: 475855-1R), washed in dH_2_O, frozen in liquid nitrogen and freeze-dried for dry mass determination and transcript analysis.

### 2.6. Phenotyping

The growth phenotype was determined on solid complete media (CM by DR. C. F. Robert (Barratt et al., 1965)), AMM and AMM supplemented with different concentrations of DES (Sigma, Art. No.: 4628) between 0.033 nM and 50 nM. Plates were inoculated with 10 µL of a 100 spores/µL spore suspension and incubated for 4 days at 37 °C in the dark. After 4 days, radial growth was measured and a picture was taken. Each growth test was done in triplicate. Additionally, the dry mass accumulation and production of SMs (i.e. Penicillin G, Sterigmatocystin, Austinol, Dehydroaustinol, Emericellamide A) were compared under the same conditions as during activation experiments (see section 2.5) with the exception of comparing DES-induced and not induced VPR4 strains.

### 2.7. SM analysis

1 mL of supernatant of *A. nidulans* liquid culture was analysed. The samples were run on a QTrap 5500 LC-MS/MS System (Applied Biosystems, Foster City, CA, USA) equipped with a TurboIonSpray electrospray ionisation (ESI) source and a 1290 Series HPLC System (Agilent, Waldbronn, Germany). Chromatographic separation was done at 25°C using a Gemini C18 150 × 4.6 mm i.d., 5 μm particle size, equipped with a C18 3 × 4 mm i.d. security guard cartridge (Phenomenex, Torrance, CA, USA). The chromatographic method and chromatographic and mass spectrometric parameters are described elsewhere (Sulyok et al., 2020). The concentration of each metabolite was then normalised to the generated dry mass during cultivation.

## 3. Results

### 3.1. Systems development and experimental design

An overview of the system and the workflow are shown in Figure 1 (plasmids) and Figure 2 (workflow). To develop the system, we consecutively transformed *A. nidulans* FGSC A1153 with two constructs. First, plasmid VPR4 (constitutively expressing *hER* and an inducible (mediated by activated hER) *VPR-dCas9*). The expression of *VPR-dCas9* can be induced by the addition of estrogens (i.e. DES) which leads to the activation of hER, its subsequent dimerisation, binding to estrogen response elements (EREs; i.e. promoter region of VPR-dCas9) and activation of respective genes (Pachlinger et al., 2005). A verified transformant carrying pVPR4 was then used as the recipient strain for the second plasmid which carries the constitutively expressed sgRNA cassette (descendants of psgRNA). The sgRNA is flanked by a hammerhead ribozyme and a HDV ribozyme to facilitate the proper processing and release of functional sgRNAs (Nodvig et al., 2015). During activation experiments, strain VPR4 was used as control strain to rule out artefacts that originate from components of plasmid VPR4 (i.e. *VPR-dCas9, hER*) or the inducer. An overview on the whole experimental workflow is shown in Figure 2.

To improve correct positioning and “landing” of sgRNA-loaded VPR-dCas9 at the selected target region, we considered two points. First, the synthetic activator should be located at a suitable distance from the expected general transcriptional machinery to ensure efficient contacts between the activator, mediator complex and the RNA-Polymerase II complex. We therefore positioned VPR-dCas9 at different distances from the predicted transcriptional start site (TSS) of the target gene. Secondly, we hypothesised that the sgRNA target sequences should not be buried in densely packed nucleosomes but optimally lie in nucleosome-free regions. We therefore performed genome-wide nucleosome accessibility assays using micrococcal nuclease digests followed by whole genome sequencing (MNase-seq). Two nucleosome positioning maps were created in the wild-type strain FGSC A4 grown at two different physiological conditions on defined AMM, i.e. conditions of active growth (i.e. primary metabolism; 15 hours shake cultures) and in stationary phase (i.e. secondary metabolism; 48 hours shake cultures). Transformants with verified correct molecular integration events of pVPR4 (i.e. strain VPR4) was analysed for SM production under non-induced and DES-induced (i.e. activating) conditions (See section 2.5) to exclude that the addition of estrogens already results in elevated levels of SMs without any sgRNA-directed targeting of VPR-dCas9 (Figure S 4). Finally, during activation experiments, sgRNA-carrying strains and control strain (i.e. VPR4) were treated exactly the same like described in section 2.5. Target gene transcription was measured by reverse transcriptase quantitative polymerase chain reaction (RT-qPCR) and the mean fold change in expression was determined by comparing sgRNA-carrying strains to control strain (i.e. VPR4). Plasmids and strains that were used or created during this study are listed in Table 1 and Table 2.

### 3.2. hER-containing strains show slight stress phenotypes in the presence of DES

As the activation system is based the estrogen-inducible *VPR-dCas9*, we first tested if the combination of hER and estrogen already led to a background activity of the SM clusters which we planned to activate at a later step. For this we first controlled the growth phenotypes of hER-containing strains and found that the activated estrogen receptor has a significant impact on growth of the strains when elevated amounts of DES are applied. The phenotype of the strain VPR4 and strains additionally carrying a guide RNA was identical. Strains that do not carry hER (i.e. FGSC A1153) are not affected by elevated DES levels (Figure S 3). To check if hER is the cause for this phenotype, two other strains that only carry the hER under a constitutive promoter were also grown under same conditions (data not shown). This experiment verified that the activated estrogen receptor is the cause for this phenotype which is in accordance with our previous observations (Pachlinger et al., 2005). The maximal inducer concentration that was tolerated by the actual strains without displaying any morphological phenotype was above 33 pM DES. 100 pM DES already showed a clear reduction in colony diameter. Consequently, for subsequent activation experiments, 33 pM DES was chosen as final inducer concentration which is also known to trigger strong activation (Pachlinger et al., 2005). Furthermore, this concentration had no effect on dry mass accumulation. The effect on SM background levels were not significant (Figure S 4). We concluded, that the system is suitable to be applied for the forced activation of the chosen SM cluster genes.

### 3.3. Activation of the silent monodictyphenone gene cluster

For a proof-of-concept, we chose the monodictyphenone (*mdp*) cluster of *A. nidulans* which was shown to be silent under standard laboratory conditions in a wild-type strain because the cluster is subject to negative chromatin-level control by the COMPASS (Bok et al, 2009) and the H3K4 demethylase KdmB (Gacek-Matthews et al., 2016). The *mdp* cluster comprises at least 10 genes. Among them are *mdpE*, which encodes a transcription factor, and *mdpG*, the backbone gene which encodes a non-reducing PKS (NR-PKS). This BGC was analysed in more detail by Chiang and colleagues who showed that forced overexpression of *mdpE* is necessary and sufficient to activate the backbone gene *mdpG* and all other cluster genes to eventually synthesise the final product monodictyphenone (Chiang et al., 2010). We therefore guided VPR-dCas9 to the *mdpE* gene promoter. An overview on the cluster, its expression profile under primary and secondary metabolic conditions and corresponding nucleosome positioning maps are shown in Figure 3.

To test the effect of different VPR-dCas9 positions in relation to the predicted nucleosome-free regions (NFRs), we designed 4 different sgRNAs according to the high-resolution map of the *mdpE* upstream region including the 5’untranslated region (UTR) and nucleosome positions as shown in Figure 4A. The sgRNAs m1 and m2 were positioned close to the predicted TSS of *mdpE*, m3 was positioned directly inside the first obvious upstream NFR which encompasses roughly 50 bps and lies around 300 bps upstream of the TSS. Finally, m4 was selected to bind even further upstream from the TSS, i.e. roughly 600 bps upstream (Figure 4A). There would be another NFR at ∼700-800 bps upstream but RNA-seq data suggest that this could very well belong to the downstream UTR of the gene AN0149 and was therefore not considered for sgRNA design. The expression analyses of *mdpE* in strains having VPR-dCas9 positioned at these different places either alone or in combination with other sgRNAs is shown in Figure 4B. Interestingly, in this promoter, positioning of VPR-dCas9 by any single sgRNA is obviously not sufficient for activation of *mdpE* because all measured transcript levels were in the range of the control strain. A clear activation potential was mediated only by sgRNA combinations of either two or all four guide RNAs. The combination of m1+m2 (m1&2) already showed a clear activation of *mdpE* in the range of 7-fold higher transcript levels compared to the background found in the control strain grown under the same conditions. When m2 was combined with m3 (m2&3) that binds to the NFR, the activation was rising to 10-fold above background levels. The highest potential was found when all four chosen sgRNAs m1 to m4 (mAll) were expressed in one strain together with VPR-dCas9. In this case *mdpE* expression was activated roughly 30-fold over the background of the control strain. These results indicate that both spatial positioning and the amount of VPR-dCas9 present at the promoter define the functionality of this fusion protein as synthetic activator.

To see if the *mdpE* expression levels are sufficiently high to yield a functional amount of the MdpE transcription factor, we tested the same samples also for expression of *mdpG*. RT-qPCR analysis shown in Figure 4C document that the forced expression of *mdpE* results in upregulation of its target gene *mdpG*. We detected a similar expression pattern of both genes. Consistent with the function of MdpE as cluster-specific TF, activation of *mdpG* was only achieved in samples where *mdpE* was expressed successfully, i.e. when at least two different sgRNAs were expressed simultaneously in the strain. Even if the expression levels of the two genes show a similar trend, it should be noted, however, that the amplitude of *mdpG* activation was much lower than for the “direct” VPR-dCas9 target *mdpE* (note the scale in y-axes of Figure 4B and Figure 4C).

### 3.4. Activation of a silent BGC of unknown function

As second example for proof-of-concept, we chose a predicted gene cluster for which no experimental evidence is available so far and no cognate metabolite is known. Genes at this target location are predicted to be part of the BGC-AN8504 (Inglis et al., 2013) which is a putative *gliP*-like non-ribosomal peptide synthetase (NRPS) (Cerqueira et al., 2014). A cluster prediction program delineates the putative BGC (Prediction tool antiSmash (Medema et al., 2011)) comprising presumably 16 genes from AN8495 to AN8508 and containing the putative NRPS-encoding gene AN8504 as well as a putative TF encoded by AN8506 (Figure 5). The nucleosome maps of this region show that the predicted BGC has a number of pronounced nucleosome free regions which made it suitable for our tests.

We targeted the promoter of the TF gene AN8506 which seems to share a very small 305 bp-control region with the divergently transcribed gene AN8507 encoding a predicted transmembrane transport protein (Cerqueira et al., 2014). The control region contains a clearly formed NFR that would perfectly serve as entry point for sgRNAs to position our activator. Interestingly, despite sharing a common promoter the transcriptional levels differed between both genes under previously tested conditions (Schinko et al., 2010, Gacek-Matthews et al., 2016; see Figure 5). While AN8506 (TF) was weakly transcribed in both 17h and 48h cultures (nutrient-limited conditions; see RNA-seq data in Figure 5), the transmembrane protein AN8507 was completely silent. This bidirectional promoter region therefore served a suitable site to test how the positioning of VPR-dCas9 in relation to the TSS of the divergently transcribed genes influences their transcription pattern. The expression profiles and nucleosome positioning maps of all genes putatively forming a BGC are shown in Figure 5.

AN8506 and AN8507 are controlled by a bidirectional promoter of only 305 bps in length that clearly contains a NFR roughly in the middle (Figure 6A). We designed two different sgRNAs for targeting VPR-dCas9 inside the NFR. The idea was to test how the position of the activator in relation to the gene would influence its activation potential. sgRNA A1 was designed to bind close to AN8506 and A2 should bind close to AN8507. The high-resolution nucleosome positioning map in Figure 6A shows that the +1 nucleosome of the moderately expressed gene AN8506 is very well positioned at both time points while this is not the case for AN8507. In the activation experiment shown in Figure 6A, we found that AN8506 is moderately activated by both sgRNAs in a similar range, i.e. roughly 6-fold by A1 and 3.5-fold by A2 suggesting that A1 positions VPR-dCas9 slightly better in relation to the general transcriptional machinery responsible for AN8506 transcription. Although the promoter architecture suggests a bidirectionally active regulatory unit, the divergently transcribed AN8507 gene is far stronger activated than AN8506 by both A1 and A2. The fact that the activation potential of VPR-dCas9 is weaker on genes that are already moderately expressed, supports this finding (Chavez et al., 2016). However, when comparing the effect of sgRNA A1 and A2 for the activation of gene AN8507, it gets apparent that even small changes in positioning of the activator leads to huge differences in the resulting gene activation. sgRNA A1 and A2 are only 50 bps apart from each other. Despite this small genomic distance, VPR-dCas9 can upregulate gene AN8507 only 30 fold when positioned by sgRNA A1 but is able to boost the transcription to 4,000 fold when positioned by sgRNA A2 (compared to control strain VPR4)! While a molecular explanation for the huge difference in activation potential remains obscure, this case documents again how significant the impact of sgRNA positioning in relation to the (predicted) TSS of the target gene is.

Next, we also wanted to see whether upregulation of the TF AN8506was sufficient to result in the expression of the NRPS-encoding gene AN8504.We therefore tested the same samples for transcription of the AN8504, but our RT-qPCR results did not detect any signal in these samples (data not shown). This indicates that expression of the TF or the transporter gene residing inside a BGC is not sufficient to turn on the backbone gene of the predicted cluster or that TF AN8506 does not positively regulate the backbone gene AN8504 at all. Using our novel VPR-dCas9 system and targeting different sgRNAs to all genes predicted to form the BGC would be a promising way to upregulate the whole cluster independently of a transcription factor. Therefore, it still could be possible to assign a product to this cluster and even to decipher the cognate biosynthetic pathway.

## 4. Discussion

Programmable transcriptional regulation is a powerful tool for the study of gene functions in general and for the activation of silent fungal BGCs in particular. The system presented here features a couple of advantages over traditional promoter replacements, *in trans* TF-overexpression or heterologous expression strategies previously used to control gene expression.

### 4.1. Keeping the genomic arrangement of the targeted locus intact

One advantage of the system is that the genetic locus of interest (e.g. a predicted BGC for which no product is known so far) is not perturbed by introduced recombinant DNA. In traditional systems, artificially introduced constitutive or inducible promoters are recombined at the locus usually with a selection marker and vector sequences (Weld et al., 2006). These changes in genomic locus arrangements may lead to silencing of the introduced genetic material (e.g. selection marker) or nearby genes if the integration event is located directly in, or adjacent to a heterochromatic region due to position effect. This was shown in *Drosophila* (position effect variegation (Elgin and Reuter, 2013; Henikoff, 1990)), yeast and also filamentous fungi (telomere position effect (Allshire et al., 1994; Gottschling et al., 1990; Palmer and Keller, 2010)). The VPR-dCas9 activator functions in the context of the native locus similar to a classical transcription activator. The sgRNA-dCas9 module functions quasi as DNA-binding-domain of this synthetic factor and by a simple change of the sgRNA sequence it can easily be targeted at different positions in relation to the core promoter (TATA or CCAAT box) (Chang et al., 2013; Kinghorn and Turner, 1992). It is well established that the interplay between the general transcription factor machinery which assembles at the core promoter as pre-initiation complex (PIC) and the transcriptional activators binding to control elements upstream of the PIC is critical for promoter activity (Soutourina, 2018). This is because the Mediator Complex must be correctly situated to be able to connect the factors bound at the control elements to the PIC at the core promoter. If this arrangement is not optimal, promoter activity will be low (Dobi and Winston, 2007). This aspect becomes particularly important when bidirectional promoters are targeted. Insertion of recombinant DNA during promoter replacement strategies disrupt co-regulation of the neighbouring genes risking regulatory off-target effects on the other gene with unknown consequences (loss- or gain-of-function phenotypes).

### 4.2. Providing an optimal landing platform and distance for the activator to the general transcription machinery

Stable interaction of an activator with DNA in promoter control elements is facilitated by binding of the factor to a NFR. Typical RNA-Pol II-transcribed fungal promoters feature such NFRs in a distance of average ∼200 bp upstream of the start codon respectively from -111 to - 5 of transcriptional start sites (TSSs)(Chen et al., 2013; Yuan et al., 2005). As the NFR is flanked by nucleosome -1 at the promoter proximal and nucleosome -2 at the promoter distal side, the location of the NFR in relation to the core promoter is clearly also critical. The possibility to guide VPR-dCas9 to different positions within a promoter is another invaluable advantage of the presented system. Of course, to be able to provide an optimal landing platform for VPR-dCas9 in the NFR of a targeted gene promoter, a nucleosome positioning map of this locus needs to be available. Optimally, recording of these nucleosome maps by MNase-Seq should be done under the same conditions than the planned VPR-dCas9 activation because it is known that some genes have NFRs that are not static and thus nucleosomes may be differently positioned under changing environmental or developmental conditions (Lai and Pugh, 2017). For example, the nitrate-responsive nucleosome shifting in the *niiA-niaD* intergenic region in *A. nidulans* is one of the paradigmatic examples for such regulation (Berger et al., 2008; Berger et al., 2006; Muro-Pastor et al., 1999). In any case, the possibility to move activator positioning sites to different positions in respect to the core promoter represents a unique advantage of the dCas9-based activators.

### 4.3. Advantages of the dCas9-based system over traditional TF-overexpression strategies

Several silent BGCs have been successfully activated by overexpression of the proprietary TF gene from a distant locus. The rationale of this approach is that a predicted TF-encoding gene lying next to genes probably forming a BGC is most likely the activator of all genes in its vicinity. Most likely, they all together would form a functional BGC, but so far nobody was able to find conditions under which these genes – including the TF - is expressed and hence the cluster product is not known. Therefore, the cluster-external forced expression of the TF may activate all genes (ideally the corresponding BGC) under its control and a novel product may be identified. However, there are two important limitations to this approach: (i) many BGCs do not contain a predicted TF (Keller, 2019), and (ii) if the overexpressed TF requires condition-specific post-translational modifications (e.g. phosphorylation, protein processing, etc.) for its activity, but the corresponding conditions are not known, the TF will be present but inactive (Karin and Hunter, 1995; Mingot et al., 2001). The VPR-dCas9 system presented here can circumvent these problems as no PTMs are known to be required for its activity.

### 4.4. A possibility to stepwise activate the cluster and hence reveal the biosynthetic pathway

Although we have not yet demonstrated its feasibility in our proof-of-concept work the system provides the possibility to activate the cluster stepwise. By designing a sgRNA-multiplex system similar to the procedure published by Nødvig and colleagues (Nodvig et al., 2018), one, two or more genes within a predicted BGC can be activated simultaneously. By activating the core biosynthetic backbone-gene (PKS, NRPS, prenyl transferases, terpene cyclases or hybrids thereof) in combination with different putative modifying or decorating enzymes and transporters will provide the possibility to elucidate different metabolic products originating from the cluster. If each step is accompanied by high-resolution mass spectrometry of the resulting cultures new products and their biosynthetic intermediates can be defined and assigned to the corresponding gene. The VPR-dCas9 system thus not only provides the possibility to study the effect of different gene combinations on the metabolic profile, but at the same time the individual biosynthetic steps could be deciphered.

### 4.5. A discovery platform suitable for many genes and fungi

The transfer of this technique to many other fungal species is highly likely as long as a suitable vectors and reliable transformation protocols exists. Even for genetically less amenable fungi a gene-free delivery of the sgRNA-loaded VPR-dCas9 synthetic activator can be envisaged. It has been shown already for fungi to work for mutagenesis approaches (Foster et al., 2018; Kuivanen et al., 2019). However, only a low copy number of the Cas9 enzyme is required when Cas9 is designed for mutagenesis and loss of the nuclease and sgRNA vectors would even be an advantage after mutagenesis has been achieved. In contrast, it is likely that a much higher number of synthetic VPR-dCas9 activator molecules are required when it comes to act as efficient transcriptional regulator at one or more gene promoters. There is no experience to these approaches and future experiments will test its applicability. Application of the system obviously is not restricted to BGCs. It is likely to work as a shuttle system for chromatin and DNA modifying enzymes altering nucleosomal PTM signatures and the chromatin architecture at the targeted locus. dCas9-fusions to such chromatin or DNA modifiers are thus a promising tool to study downstream pathways involved in epigenetic regulation. It should also be mentioned that dCas9 could fused to a transcriptional repressor or for transcriptional interference (CRISPRi), both inactivating genes of interest, e.g. in knock-down approaches for essential genes (Hilton et al., 2015; Larson et al., 2013).

### 4.6. Guide RNA design as a key factor for activation

The design and positioning of the sgRNAs (i.e. position of VPR-dCas9 in the genome) is a crucial factor for proper function. The majority of Cas9 applications concern the cutting of one or both DNA strands within a gene body (Ran et al., 2013). Once the Cas9 protein can cause a nick or a double strand brake that is erroneously repaired by the DNA repair machinery, presence of Cas9 at the locus is neither necessary nor likely anymore (Brinkman et al., 2018). The story is however different for the activator VPR-dCas9. Not only is the protein bigger, which can lead to steric hindrances at the desired locus, but the outcome of an experiment is not determined by a one-time event (i.e. erroneous DNA repair with Cas9) but is the result of a persistent presence of the activator at the desired locus. The residence time of dCas9 at the target locus also greatly depends on the guide RNA (Ma et al., 2016). Many transcriptionally relevant proteins and their binding sites (e.g. RNA-Pol II complex, nucleosomes, Transcription factors, CCAAT box binding proteins, TATA box, etc.) are in the promoter which on the one hand should not be blocked but on the other hand, can interfere with the VPR-dCas9 activator (Chen et al., 2013; Mao and Chen, 2019; Nikolov and Burley, 1997). Furthermore, the activator has to be positioned at a location where it can actually trigger transcriptional activation. However, if the sgRNA is designed in a way which positions VPR-dCas9 too close to the TSS, the transcription could get blocked or reduced due to CRISPRi effects (Larson et al., 2013). Therefore, the chromatin and other proteins at the target location can have a major impact on accessibility of VPR-dCas9 to the DNA and therefore its efficiency (Chen et al., 2016; Chung et al., 2020). Many promoters contain loosely attached nucleosomes or even NFRs which could be good entry points for VPR-dCas9 (Yadon et al., 2010).

### 4.7. Limitations of the system (as tested so far)

Despite our promising first results and hypothetical great possibilities, we observed during our work several important limitations and possible hurdles for an establishment of the system as efficient expression platform. The shortcomings we experienced were e.g. quite strong variation in expression strengths. For example, the activation pattern of *mdpG* is similar to the induction of *mdpE* but overall expression is much weaker. There is no obvious explanation for that. However, one possibility could be that *mdpA* - an important coactivator for the monodictyphenone BGC (Chiang et al., 2010) - was not targeted and activated together with *mdpE*. It was shown that the cluster activity is significantly reduced if *mdpA* is deleted. However, it is still not clear if *mdpA* is under the transcriptional control of MdpE. It could also be that they are regulated independently and that complex formation of MdpE and MdpA is needed to exert its full activating function on the other genes of the cluster. Such a scenario would be consistent with our results. In addition, the strength of activation depends on the position of the gRNA. In case of the very short bidirectional promoter between genes AN8506 and AN8507 even a distance of 50 bps can change the activation potential for about a thousand-fold. It is unlikely that only the short distance accounts for the difference in activation. It seems more likely that other (regulatory) proteins or protein complexes are either hindered during assembly by VPR-dCas9 or, *vice versa*, that the activator is not able to access the DNA properly. To know the exact proteomic composition at the promoter would help evaluating the outcome of the experiments and give more insight about efficient positioning of gRNAs for optimal gene activation.

## 5. Summary

The fact that activations that range in the thousand folds compared to the basal, non-induced level can be achieved makes it a very promising tool for forced gene activation approaches. To effectively use this tool, the proper design of guide RNAs has to be further studied and optimised. The finding of patterns for *in silico* prediction of best sgRNA binding sites would demand in depth research on more promoter/gene types with larger experimental setups (“guide RNA walking” along the promoter) and detailed promoter characterisation like local proteomic composition analyses at the targeted promoters.

## Acknowledgements

We want to thank colleagues from the Perrimon laboratory (Harvard Medical School, Boston, Massachusetts, USA) for sending us the vectors for the amplification of the VPR gene, as well as colleagues from the Leonardt laboratory (LMU Munich, Martinsried, Germany) who sent us the plasmids for the amplification of dCas9, and from the Mortensen laboratory (Technical University of Denmark, Søltofts Plads, Kongens Lyngby, Denmark) that provided the templates of the sgRNA cassette. We are also thankful to Matthias Steiger and Michael Sauer (BOKU University) for help with the *coxA* promoter design. Many thanks also to Dragana Bandian (AIT, Austrian Institute of Technology) who provided the gDNAs of fungal wildtype isolates for plasmid construction. Work was funded by grant Nr. K3-G-2/026-2013 “Bioactive Microbial Metabolites” from the NFB Lower Austria Science Fund and grant P32790 “ChroCosm” from the FWF Austrian Science Fund to JS and by the “Wirtschaftskammerpreis 2018” from the WKO Wien to AS.

## Supplemental Material

**Figure S 1:**
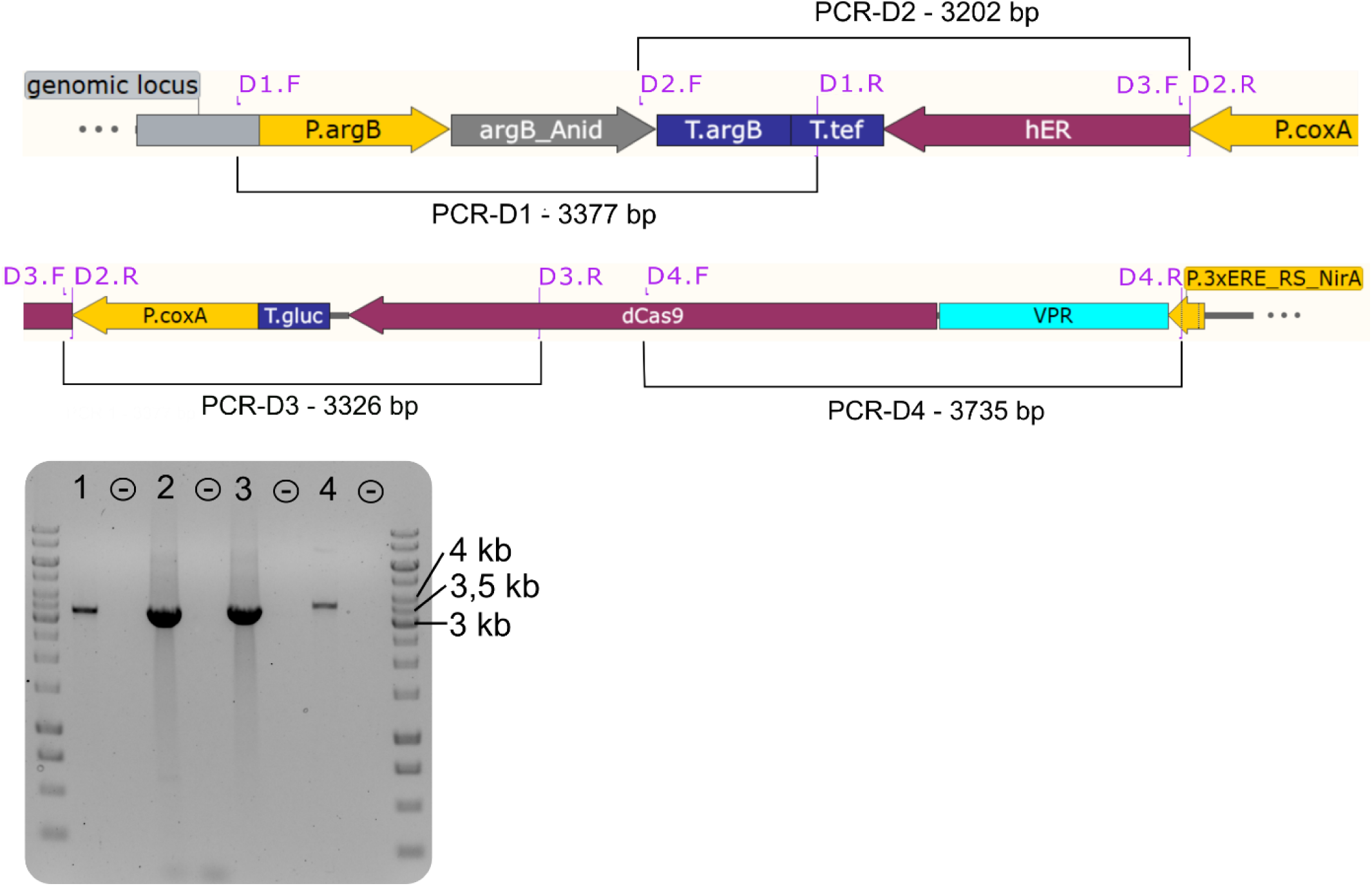
Diagnostic PCRs of Strain VPR4. The 5’ part of plasmid VPR4 is depicted as it would integrate into the desired genomic locus (*argB*). The “genomic locus” at the 5’ of the construct (in grey) is the upstream region of the *argB* gene that is not present on the plasmid VPR4. Primer pairs (violet) (Dx.F and Dx.R (x=1-4))) are connected by a black lines which correspond to the PCR product (D1-D4). The results of the diagnostic PCR for Strain VPR4 can be retrieved from the gel picture where each amplicon (1= D1, 2= D2, etc.) is followed by a negative control (-). Hence, Strain VPR4 has plasmid VPR4 in the desired locus (*argB*) and possesses all necessary features for the expression of the activator VPR-dCas9. See supplementary Table S 1 and Figure 1 for more detailed information about primers and the plasmid respectively.

**Figure S 2:**
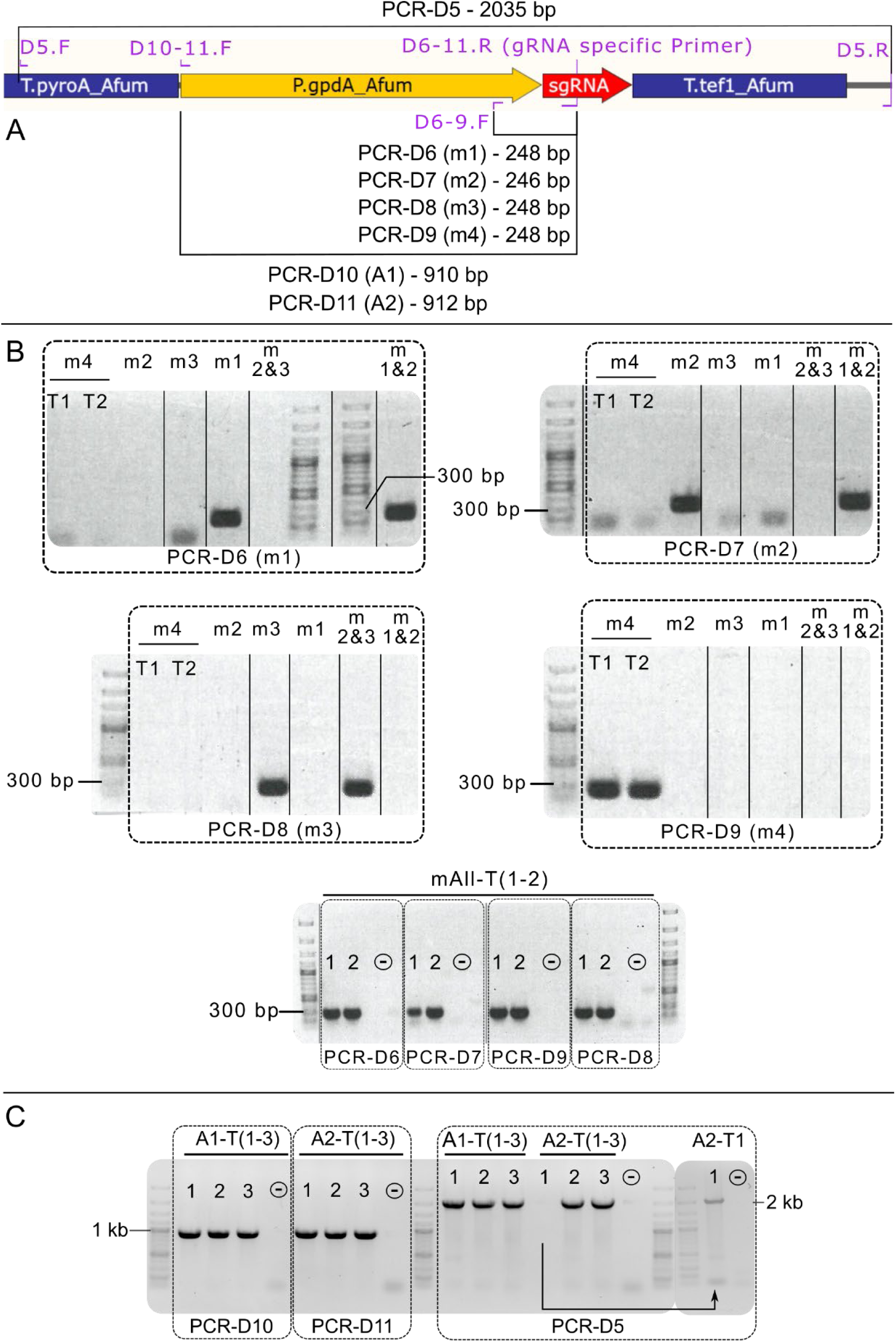
Diagnostic PCRs of single guide RNA carrying strains. **A.** Section of the plasmid map containing the sgRNA cassette. Primer pairs (violet) (Dx.F and Dx.R (x=5-11)) are connected by a black line which correspond to the PCR product (D5-D11). The results of the diagnostic PCR for Strain VPR4 can be retrieved from the gel pictures in **B** (activation strains for *mdpE*) and **C** (activation strains for AN8506 and AN8507). The strain description was abbreviated by omitting “VPR4-” (i.e. A1-T1 corresponds to VPR4-A1-T1). The abbreviated strain names are displayed above the respective lanes. Each box (dashed lines) represents the results of a primer pair (bottom side of the box). Solid lines separating lanes indicate that dispensable lanes of the gel picture were removed. Information about the primer pairs are included in Table S 1

**Figure S 3:**
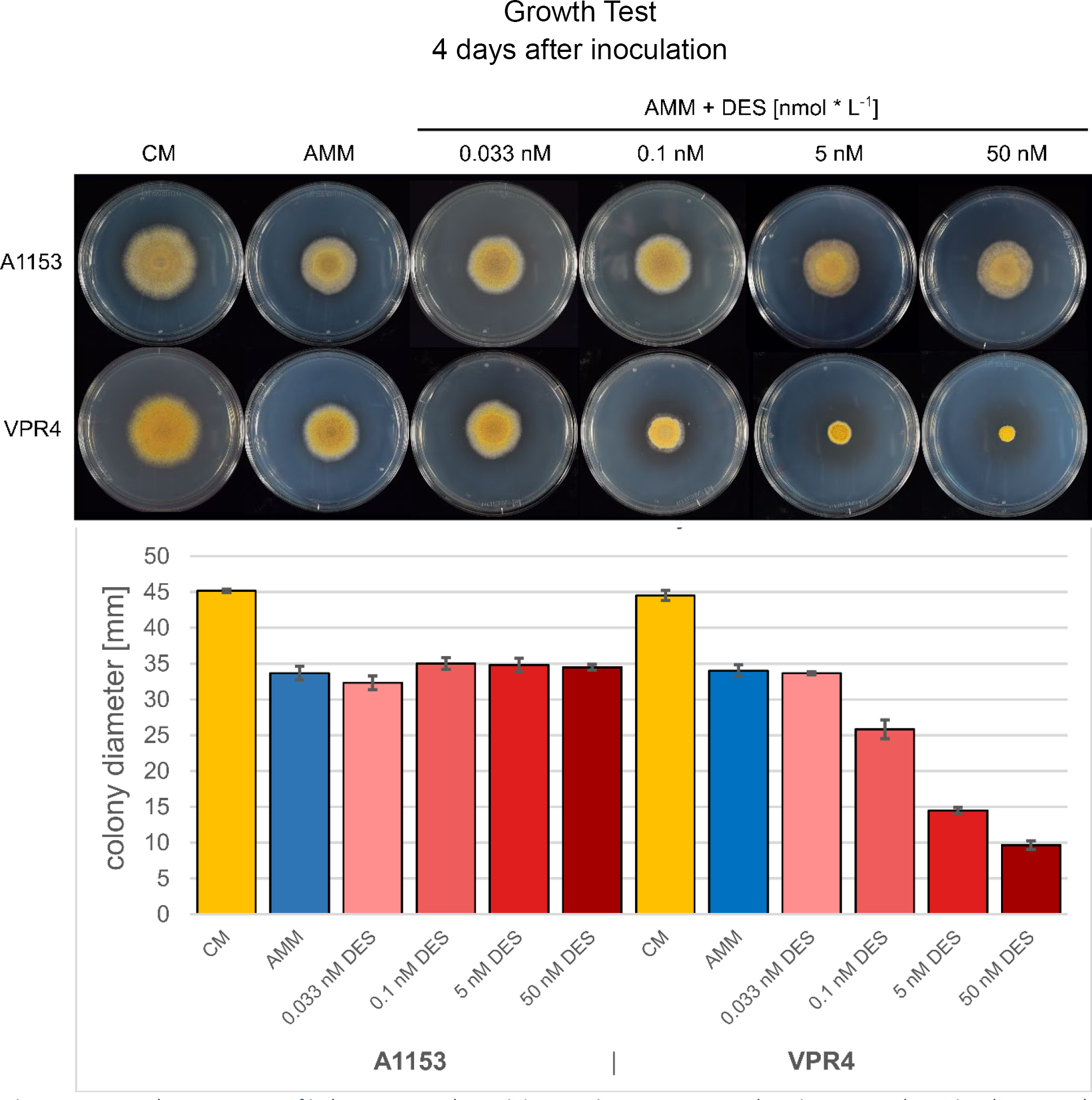
Growth Test. Impact of inducer DES on the recipient strain FGSC A1153 and strain VPR4. Only strains that carry the hER are affected by the inducer DES. The recipient strain does not show any phenotypical changes upon incubation with DES. Activation of the estrogen receptor (Strains VPR4 and descendants) leads to growth phenotypes. At 33 pM DES (concentration for activation experiments), no impact on radial growth or morphology can be observed. At 100 pM DES, a significant radial growth reduction of ∼ 40% can be observed. At 50 nM DES, the fungus does not pass the inoculation radius but still can sporulate (radial growth reduction of 100%). Inoculations were done in triplicate and growth was measured after a time period of 4 days

**Figure S 4:**
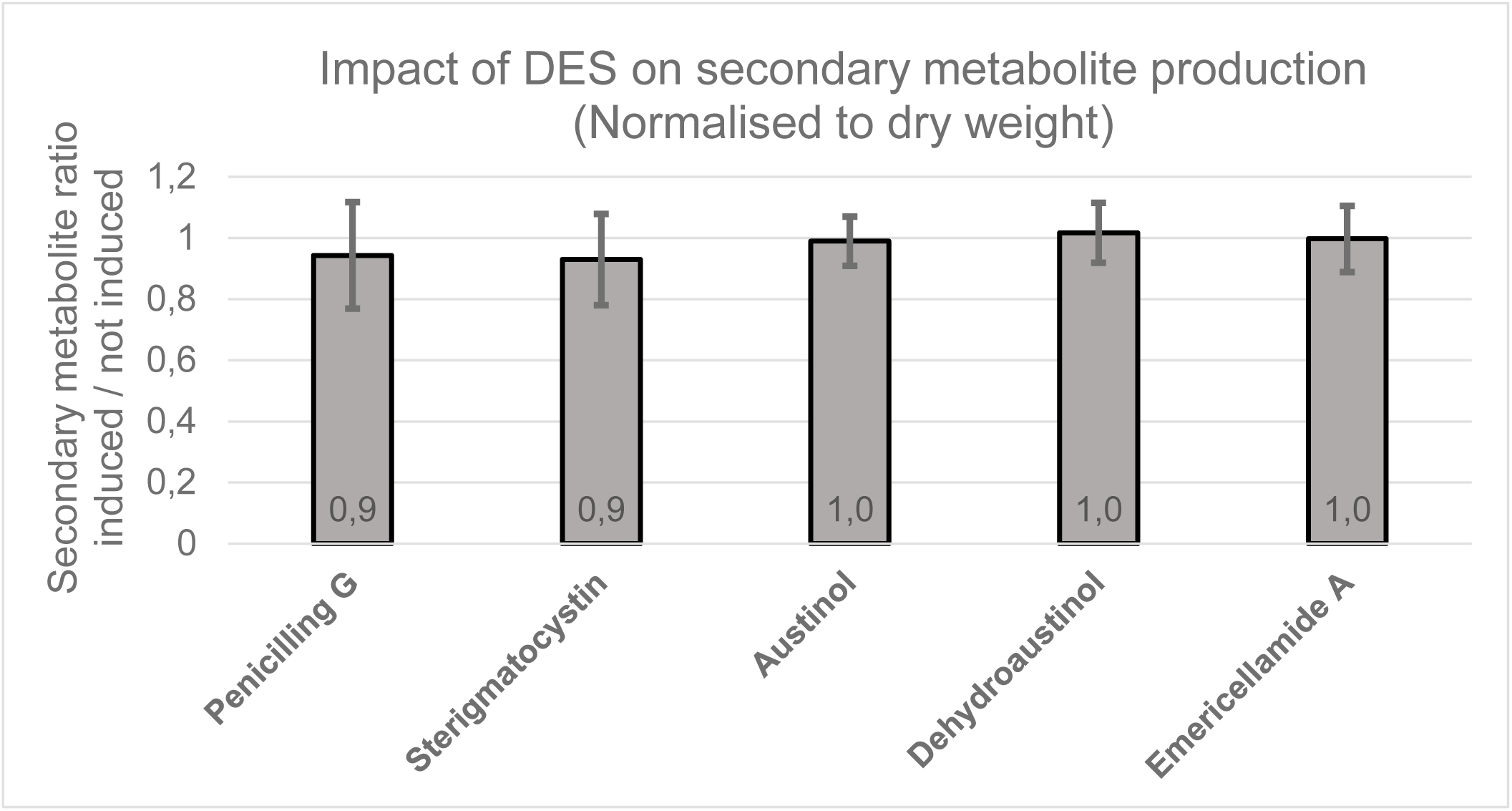
Impact of DES addition on the production of SMs in a hER carrying strain. Supernatants for chemical analysis of *A. nidulans* SMs were taken 8 hours after induction (i.e. 48 hours total inoculation time). SM production was normalized to dry weight accumulation which was not influenced by DES addition. SM production was calculated by dividing the SM levels in the DES-induced by the non-induced state. All strains carry the *hER* gene and the fusion gene *VPR-dCas9*. The dataset comprises 3 different experiments with a total sample count of 22 (i.e. 11 times induced and 11 times not induced) The SM profile shows no significant change.

**Table S 1:**
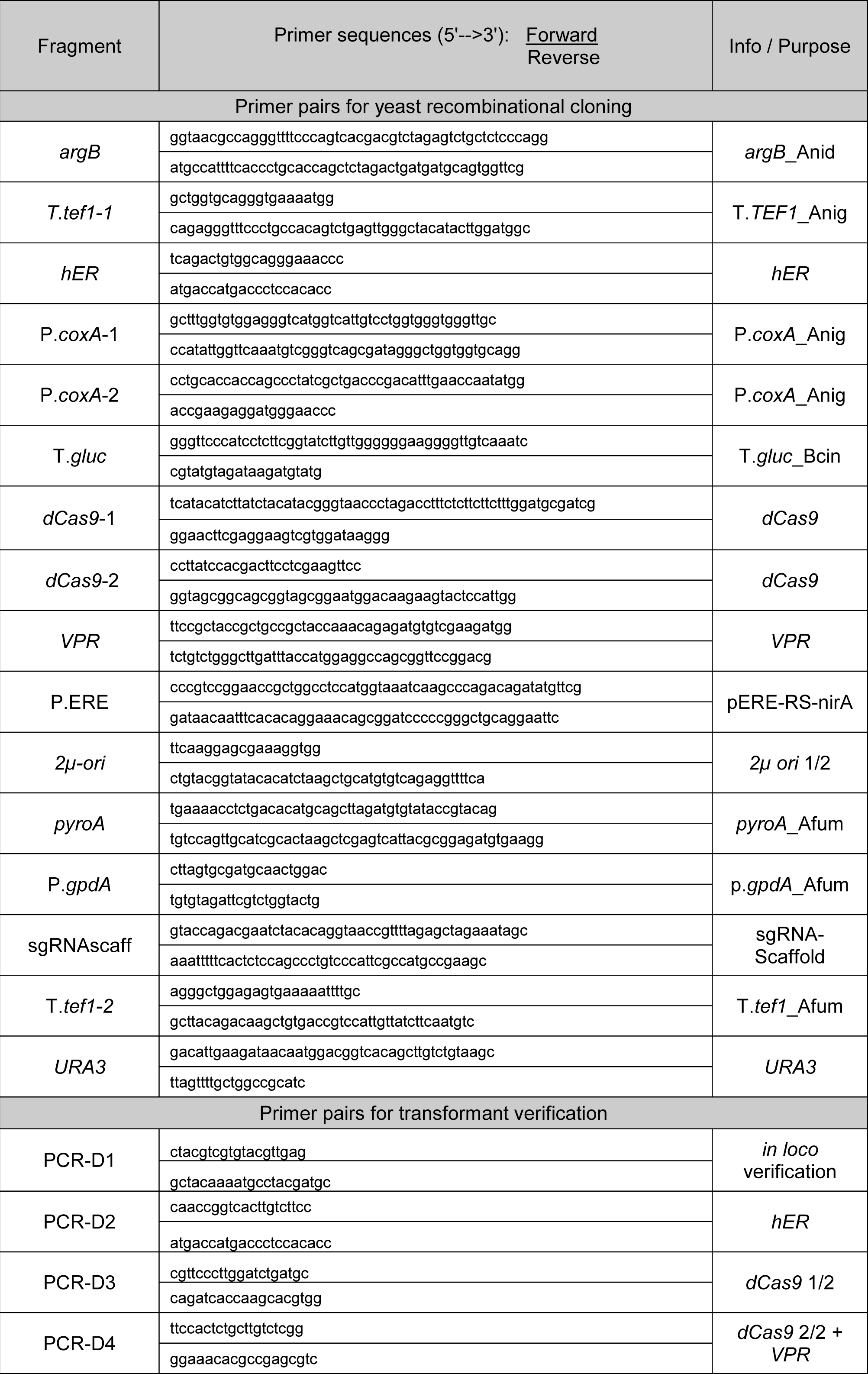
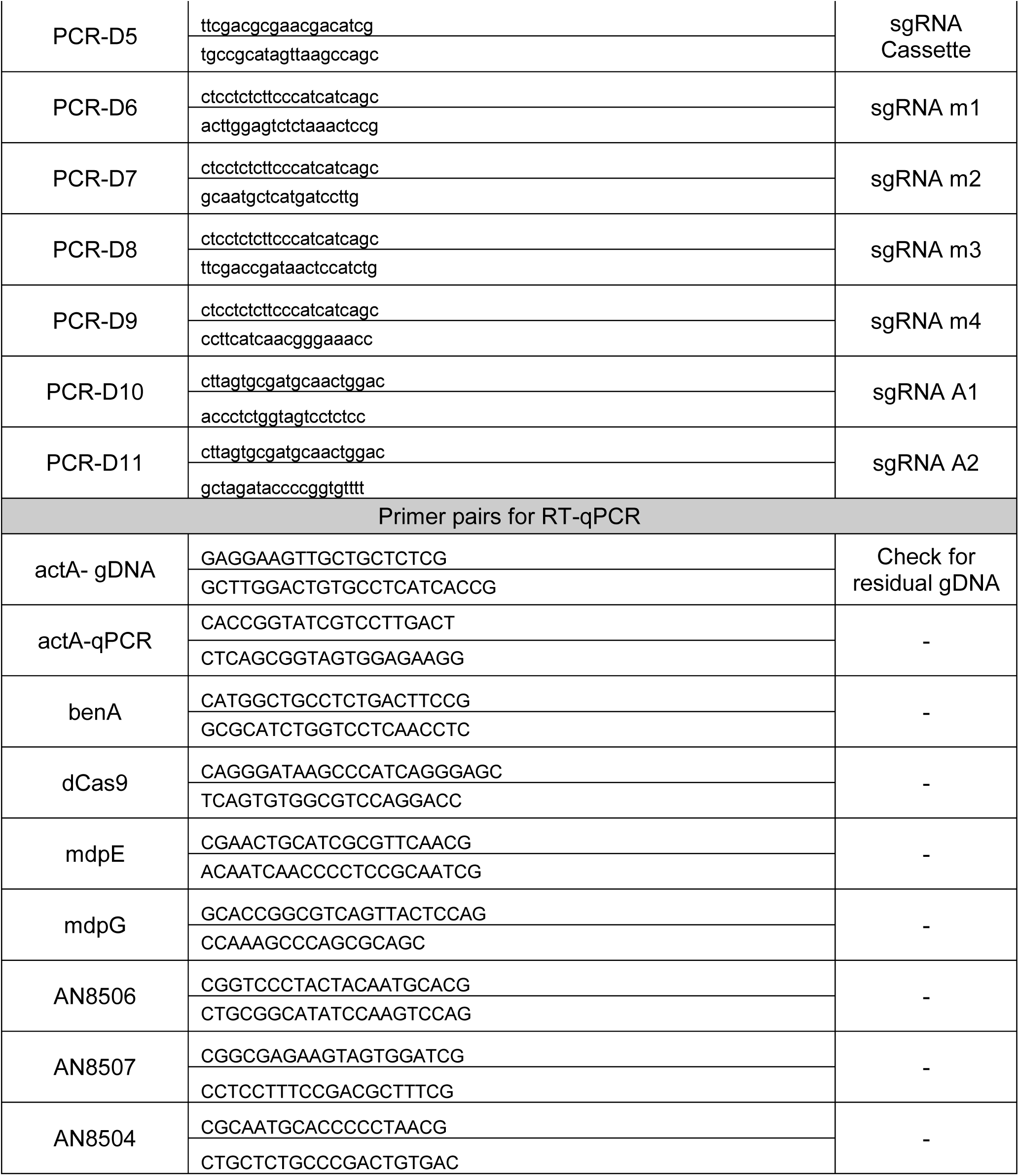
<u>Primersets</u> used during this study

**Table S 2:**
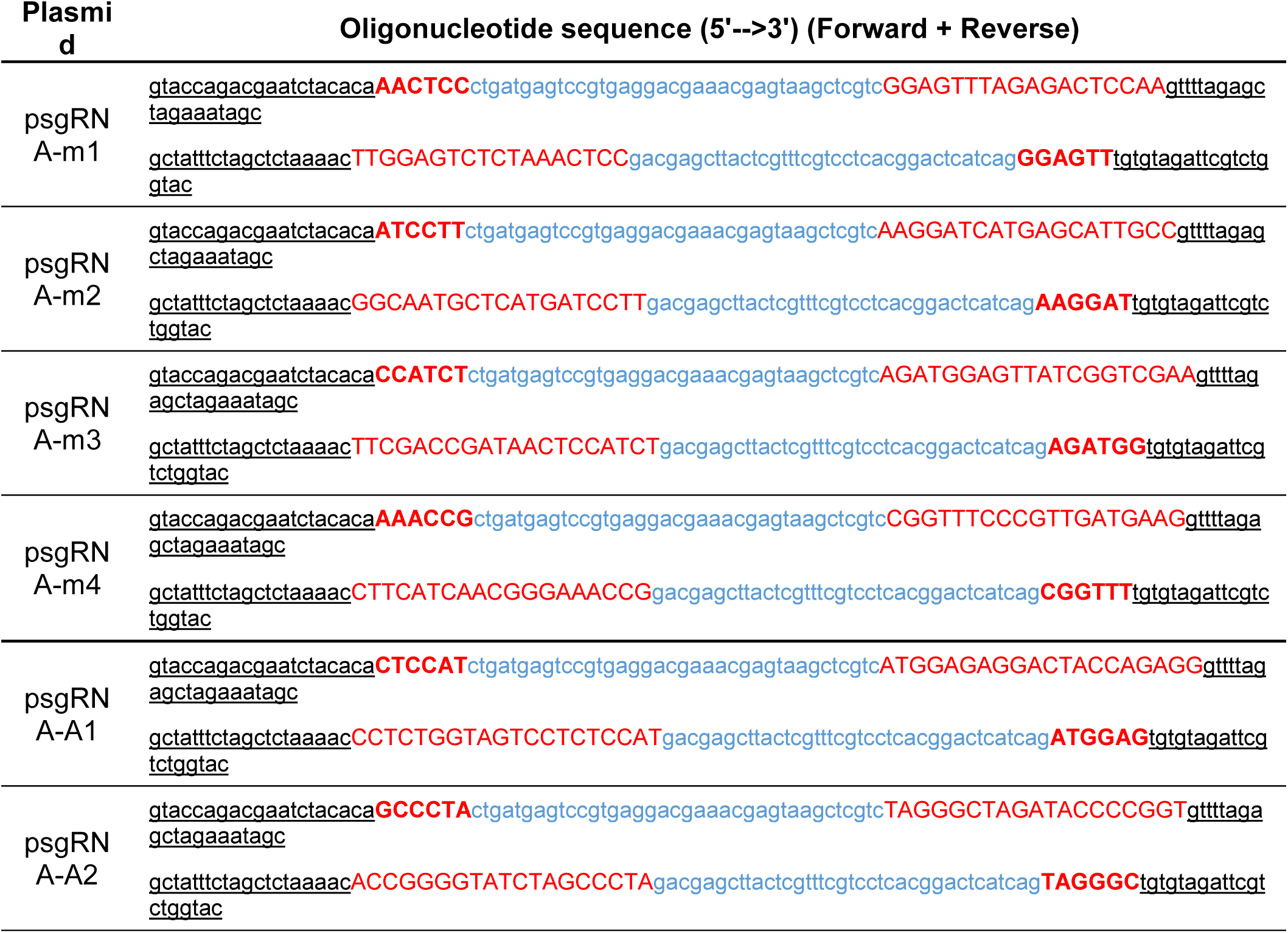
<u>Oligonucleotides for cloning of sgRNA carrying plasmids</u> (descendants of psgRNA). Forward and reverse oligonucleotide consist of overhangs (black, underlined letters; needed for YRC), inverted repeat of 5’ protospacer (in red, bold letters needed for proper Hammerhead ribozyme function), Hammerhead motif (blue letters) and the protospacer sequence (red letters; i.e. target locus specific nucleotide sequence). Prior to YRC, the oligonucleotides had to be annealed as described in section 2.1.

