## Supplemental material for "Activation of silent secondary metabolite gene clusters by nucleosome map-guided positioning of the synthetic transcription factor VPR-dCas9"

### Supplemental Materials

Andreas Schüller[1], Lisa Wolansky[1], Harald Berger[1], Lena Studt[1], Agnieszka Gacek-Matthews[1§], Michael Sulyok[2], and Joseph Strauss[1*]

[1]Institute of Microbial Genetics, Department of Applied Genetics and Cell Biology, [2] Institute of Bioanalytics and Agrometabolomics, Department of Agrobiotechnology, BOKU-University of Natural Resources and Life Sciences Vienna, BOKU-Campus Tulln, Tulln/Donau, Austria.

[§]present address: Institute of Microbiology, Functional Microbiology Division, University of Veterinary Sciences Vienna.

Address: Joseph Strauss, Fungal Genetics Lab, Institute of Microbial Genetics, Department of Applied Genetics and Cell Biology, BOKU-University of Natural Resources and Life Sciences Vienna, BOKU-Campus Tulln, Konrad Lorenz Strasse 24, A-3430 Tulln/Donau, Austria. Tel. +43-147654-94420

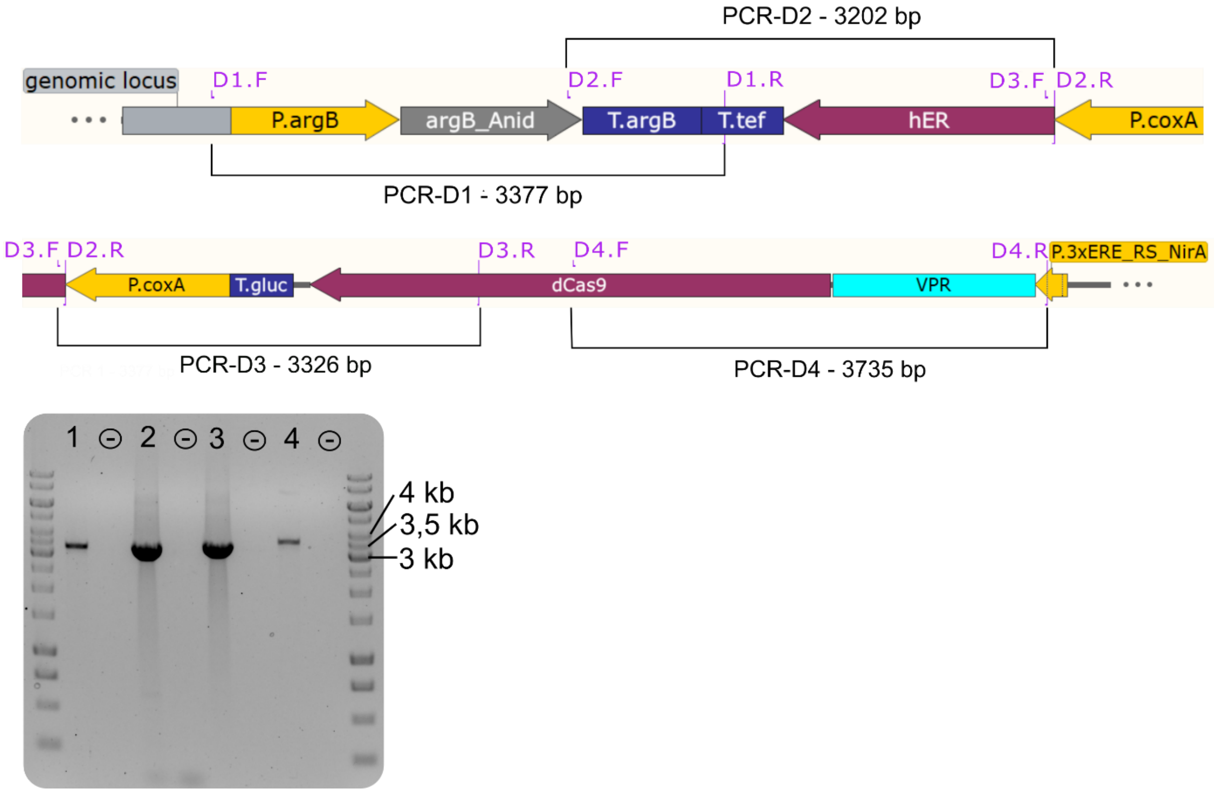


Figure S 1: Diagnostic PCRs of Strain VPR4. The 5' part of plasmid VPR4 is depicted as it would integrate into the desired genomic locus (*argB*). The "genomic locus" at the 5' of the construct (in grey) is the upstream region of the *argB* gene that is not present on the plasmid VPR4. Primer pairs (violet) (Dx.F and Dx.R (x=1-4))) are connected by a black lines which correspond to the PCR product (D1- D4). The results of the diagnostic PCR for Strain VPR4 can be retrieved from the gel picture where each amplicon (1= D1, 2= D2, etc.) is followed by a negative control (-). Hence, Strain VPR4 has plasmid VPR4 in the desired locus (*argB*) and possesses all necessary features for the expression of the activator VPR-dCas9. See supplementary Table S 1 and **Error! Reference source not found.** for more detailed information about primers and the plasmid respectively.

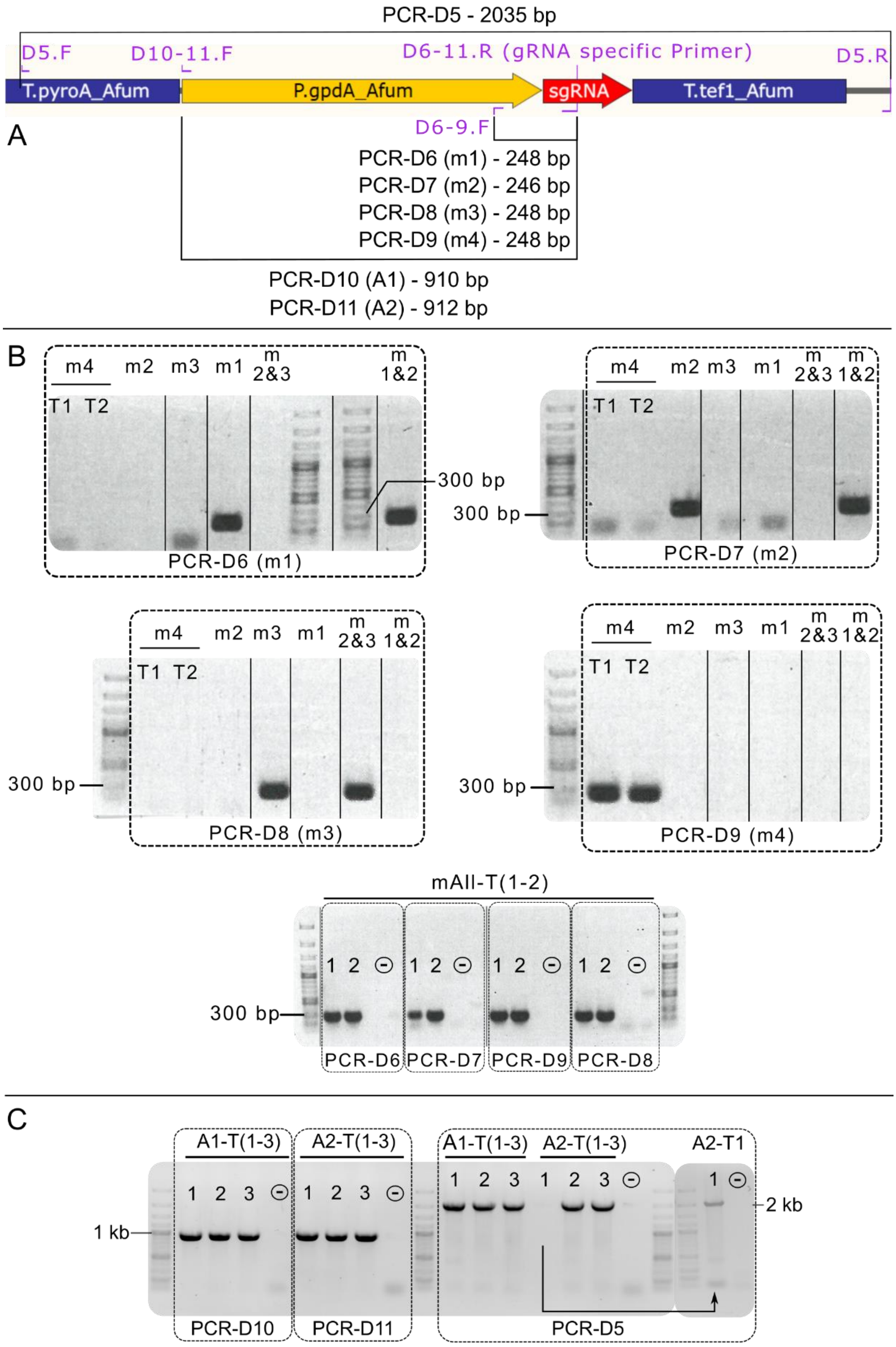


Figure S 2: Diagnostic PCRs of single guide RNA carrying strains. **A.** Section of the plasmid map containing the sgRNA cassette. Primer pairs (violet) (Dx.F and Dx.R (x=5-11)) are connected by a black line which correspond to the PCR product (D5- D11). The results of the diagnostic PCR for Strain VPR4 can be retrieved from the gel pictures in **B** (activation strains for *mdpE*) and **C** (activation strains for AN8506 and AN8507). The strain description was abbreviated by omitting "VPR4-" (i.e. A1-T1 corresponds to VPR4-A1-T1). The abbreviated strain names are displayed above the respective lanes. Each box (dashed lines) represents the results of a primer pair (bottom side of the box). Solid lines separating lanes indicate that dispensable lanes of the gel picture were removed. Information about the primer pairs are included in Table S 1

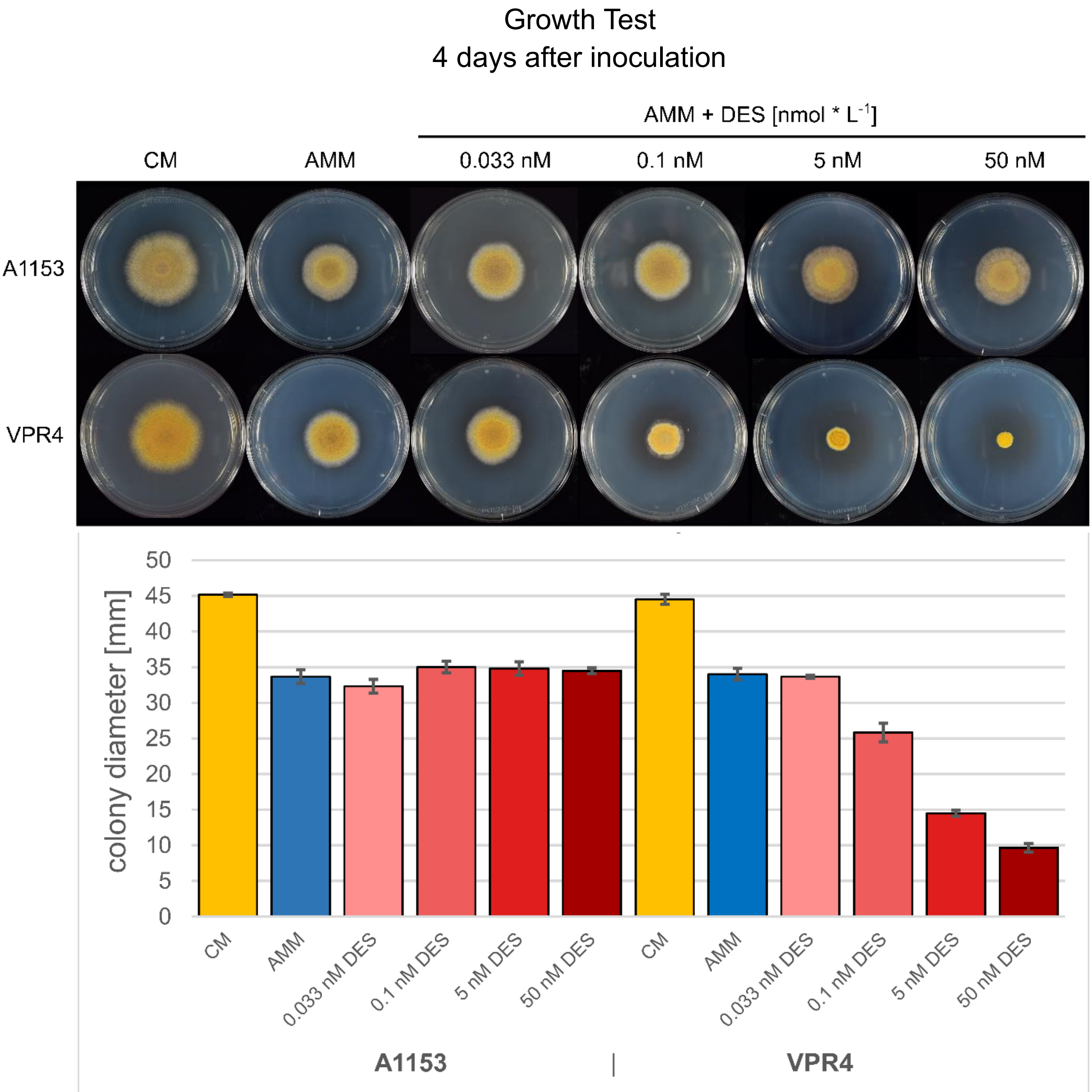


Figure S 3: Growth Test. Impact of inducer DES on the recipient strain FGSC A1153 and strain VPR4. Only strains that carry the hER are affected by the inducer DES. The recipient strain does not show any phenotypical changes upon incubation with DES. Activation of the estrogen receptor (Strains VPR4 and descendants) leads to growth phenotypes. At 33 pM DES (concentration for activation experiments), no impact on radial growth or morphology can be observed. At 100 pM DES, a significant radial growth reduction of ~ 40% can be observed. At 50 nM DES, the fungus does not pass the inoculation radius but still can sporulate (radial growth reduction of 100%). Inoculations were done in triplicate and growth was measured after a time period of 4 days

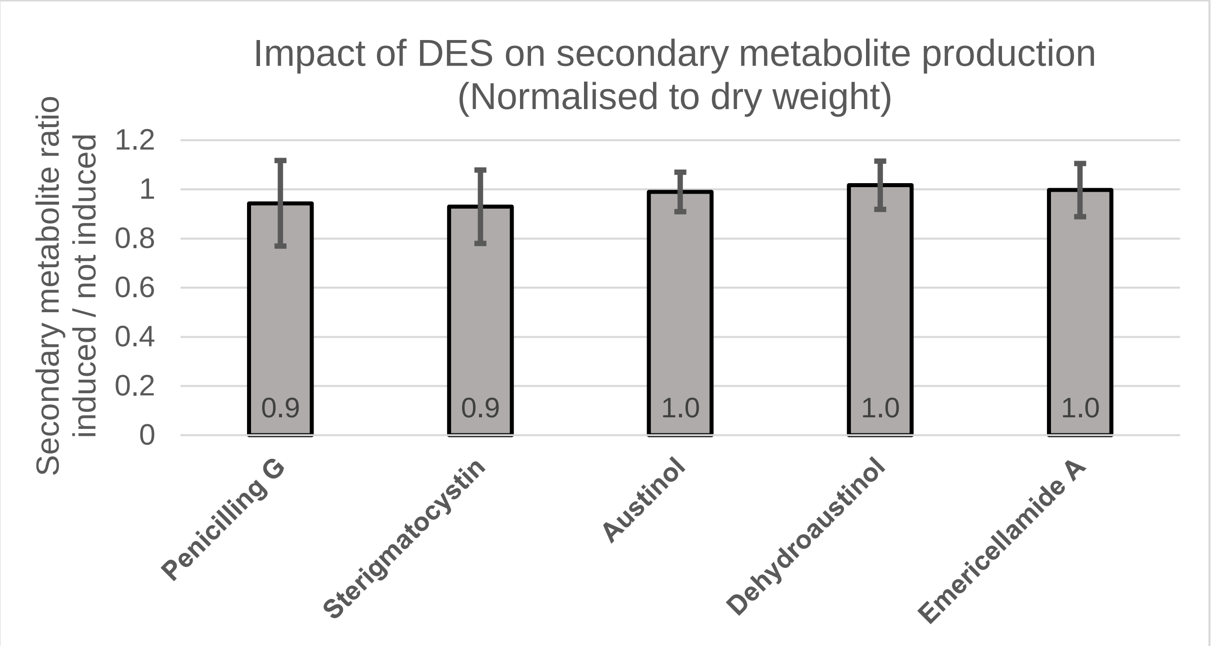


Figure S 4: Impact of DES addition on the production of SMs in a hER carrying strain. Supernatants for chemical analysis of *A. nidulans* SMs were taken 8 hours after induction (i.e. 48 hours total inoculation time). SM production was normalized to dry weight accumulation which was not influenced by DES addition. SM production was calculated by dividing the SM levels in the DES-induced by the non-induced state. All strains carry the *hER* gene and the fusion gene *VPR-dCas9*. The dataset comprises 3 different experiments with a total sample count of 22 (i.e. 11 times induced and 11 times not induced) The SM profile shows no significant change.

| Fragment | Primer sequences (5'-->3'): <u>Forward</u> / Reverse | Info / Purpose |
|---|---|---|
| Primer pairs for yeast recombinational cloning | | |
| *argB* | ggtaacgccagggttttcccagtcacgacgtctagagtctgctctcccagg | *argB*_Anid |
| | atgccattttcaccctgcaccagctctagactgatgatgcagtggttcg | |
| *T.tef1-1* | gctggtgcagggtgaaaatgg | T.*TEF1*_Anig |
| | cagagggtttccctgccacagtctgagttgggctacatacttggatggc | |
| *hER* | tcagactgtggcagggaaaccc | *hER* |
| | atgaccatgaccctccacacc | |
| P.*coxA*-1 | gctttggtgtggagggtcatggtcattgtcctggtgggtgggttgc | P.*coxA*_Anig |
| | ccatattggttcaaatgtcgggtcagcgatagggctggtggtgcagg | |
| P.*coxA*-2 | cctgcaccaccagccctatcgctgacccgacatttgaaccaatatgg | P.*coxA*_Anig |
| | accgaagaggatgggaaccc | |
| T.*gluc* | gggttcccatcctcttcggtatcttgttgggggaaggggttgtcaaatc | T.*gluc*_Bcin |
| | cgtatgtagataagatgtatg | |
| *dCas9*-1 | tcatacatcttatctacatacgggtaaccctagacctttctcttcttctttggatgcgatcg | *dCas9* |
| | ggaacttcgaggaagtcgtggataaggg | |
| *dCas9*-2 | ccttatccacgacttcctcgaagttcc | *dCas9* |
| | ggtagcggcagcggtagcggaatggacaagaagtactccattgg | |
| *VPR* | ttccgctaccgctgccgctaccaaacagagatgtgtcgaagatgg | *VPR* |
| | tctgtctgggcttgatttaccatggaggccagcggttccggacg | |
| P.ERE | cccgtccggaaccgctggcctccatggtaaatcaagcccagacagatatgttcg | pERE-RS-nirA |
| | gataacaatttcacacaggaaacagcggatcccccgggctgcaggaattc | |
| *2µ-ori* | ttcaaggagcgaaaggtgg | *2µ ori* 1/2 |
| | ctgtacggtatacacatctaagctgcatgtgtcagaggttttca | |
| *pyroA* | tgaaaacctctgacacatgcagcttagatgtgtataccgtacag | *pyroA*_Afum |
| | tgtccagttgcatcgcactaagctcgagtcattacgcggagatgtgaagg | |
| P.*gpdA* | cttagtgcgatgcaactggac | p.*gpdA*_Afum |
| | tgtgtagattcgtctggtactg | |
| sgRNAscaff | gtaccagacgaatctacacaggtaaccgttttagagctagaaatagc | sgRNA-Scaffold |
| | aaatttttcactctccagccctgtcccattcgccatgccgaagc | |
| T.*tef1-2* | agggctggagagtgaaaaattttgc | T.*tef1*_Afum |
| | gcttacagacaagctgtgaccgtccattgttatcttcaatgtc | |
| *URA3* | gacattgaagataacaatggacggtcacagcttgtctgtaagc | *URA3* |
| | ttagttttgctggccgcatc | |
| Primer pairs for transformant verification | | |
| PCR-D1 | ctacgtcgtgtacgttgag | *in loco* verification |
| | gctacaaaatgcctacgatgc | |
| PCR-D2 | caaccggtcacttgtcttcc | *hER* |
| | atgaccatgaccctccacacc | |
| PCR-D3 | cgttcccttggatctgatgc | *dCas9* 1/2 |
| | cagatcaccaagcacgtgg | |
| PCR-D4 | ttccactctgcttgtctcgg | *dCas9* 2/2 + *VPR* |
| | ggaaacacgccgagcgtc | |

| | | |
|---|---|---|
| PCR-D5 | ttcgacgcgaacgacatcg<br>tgccgcatagttaagccagc | sgRNA Cassette |
| PCR-D6 | ctcctctcttcccatcatcagc<br>acttggagtctctaaactccg | sgRNA m1 |
| PCR-D7 | ctcctctcttcccatcatcagc<br>gcaatgctcatgatccttg | sgRNA m2 |
| PCR-D8 | ctcctctcttcccatcatcagc<br>ttcgaccgataactccatctg | sgRNA m3 |
| PCR-D9 | ctcctctcttcccatcatcagc<br>ccttcatcaacgggaaacc | sgRNA m4 |
| PCR-D10 | cttagtgcgatgcaactggac<br>accctctggtagtcctctcc | sgRNA A1 |
| PCR-D11 | cttagtgcgatgcaactggac<br>gctagataccccggtgtttt | sgRNA A2 |
| Primer pairs for RT-qPCR | | |
| actA- gDNA | GAGGAAGTTGCTGCTCTCG<br>GCTTGGACTGTGCCTCATCACCG | Check for residual gDNA |
| actA-qPCR | CACCGGTATCGTCCTTGACT<br>CTCAGCGGTAGTGGAGAAGG | - |
| benA | CATGGCTGCCTCTGACTTCCG<br>GCGCATCTGGTCCTCAACCTC | - |
| dCas9 | CAGGGATAAGCCCATCAGGGAGC<br>TCAGTGTGGCGTCCAGGACC | - |
| mdpE | CGAACTGCATCGCGTTCAACG<br>ACAATCAACCCCTCCGCAATCG | - |
| mdpG | GCACCGGCGTCAGTTACTCCAG<br>CCAAAGCCCAGCGCAGC | - |
| AN8506 | CGGTCCCTACTACAATGCACG<br>CTGCGGCATATCCAAGTCCAG | - |
| AN8507 | CGGCGAGAAGTAGTGGATCG<br>CCTCCTTTCCGACGCTTTCG | - |
| AN8504 | CGCAATGCACCCCCTAACG<br>CTGCTCTGCCCGACTGTGAC | - |

Table S 1: Primersets used during this study

| Plasmid | Oligonucleotide sequence (5'-->3') (Forward + Reverse) |
| --- | --- |
| psgRNA -m1 | gtaccagacgaatctacaca**AACTCC**ctgatgagtccgtgaggacgaaacgagtaagctcgtcGGAGTTTAGAGACTCCAAgttttagagctagaaatagc<br>gctatttctagctctaaaacTTGGAGTCTCTAAACTCCgacgagcttactcgtttcgtcctcacggactcatcag**GGAGTT**tgtgtagattcgtctggtac |
| psgRNA -m2 | gtaccagacgaatctacaca**ATCCTT**ctgatgagtccgtgaggacgaaacgagtaagctcgtcAAGGATCATGAGCATTGCCgttttagagctagaaatagc<br>gctatttctagctctaaaacGGCAATGCTCATGATCCTTgacgagcttactcgtttcgtcctcacggactcatcag**AAGGAT**tgtgtagattcgtctggtac |
| psgRNA -m3 | gtaccagacgaatctacaca**CCATCT**ctgatgagtccgtgaggacgaaacgagtaagctcgtcAGATGGAGTTATCGGTCGAAgttttagagctagaaatagc<br>gctatttctagctctaaaacTTCGACCGATAACTCCATCTgacgagcttactcgtttcgtcctcacggactcatcag**AGATGG**tgtgtagattcgtctggtac |
| psgRNA -m4 | gtaccagacgaatctacaca**AAACCG**ctgatgagtccgtgaggacgaaacgagtaagctcgtcCGGTTTCCCGTTGATGAAGgttttagagctagaaatagc<br>gctatttctagctctaaaacCTTCATCAACGGGAAACCGgacgagcttactcgtttcgtcctcacggactcatcag**CGGTTT**tgtgtagattcgtctggtac |
| psgRNA -A1 | gtaccagacgaatctacaca**CTCCAT**ctgatgagtccgtgaggacgaaacgagtaagctcgtcATGGAGAGGACTACCAGAGGgttttagagctagaaatagc<br>gctatttctagctctaaaacCCTCTGGTAGTCCTCTCCATgacgagcttactcgtttcgtcctcacggactcatcag**ATGGAG**tgtgtagattcgtctggtac |
| psgRNA -A2 | gtaccagacgaatctacaca**GCCCTA**ctgatgagtccgtgaggacgaaacgagtaagctcgtcTAGGGCTAGATACCCCGGTgttttagagctagaaatagc<br>gctatttctagctctaaaacACCGGGGTATCTAGCCCTAgacgagcttactcgtttcgtcctcacggactcatcag**TAGGGC**tgtgtagattcgtctggtac |

Table S 2: Oligonucleotides for cloning of sgRNA carrying plasmids (descendants of psgRNA). Forward and reverse oligonucleotide consist of overhangs (black, underlined letters; needed for YRC), inverted repeat of 5’ protospacer (in red, bold letters needed for proper Hammerhead ribozyme function), Hammerhead motif (blue letters) and the protospacer sequence (red letters; i.e. target locus specific nucleotide sequence). Prior to YRC, the oligonucleotides had to be annealed as described in section **Error! Reference source not found.**.
